# Benchmarking long-read RNA-sequencing analysis tools using *in silico* mixtures

**DOI:** 10.1101/2022.07.22.501076

**Authors:** Xueyi Dong, Mei R. M. Du, Quentin Gouil, Luyi Tian, Jafar S. Jabbari, Rory Bowden, Pedro L. Baldoni, Yunshun Chen, Gordon K. Smyth, Shanika L. Amarasinghe, Charity W. Law, Matthew E. Ritchie

## Abstract

The current lack of benchmark datasets with inbuilt ground-truth makes it challenging to compare the performance of existing long-read isoform detection and differential expression analysis workflows. Here, we present a benchmark experiment using two human lung adenocarcinoma cell lines that were each profiled in triplicate together with synthetic, spliced, spike-in RNAs (“sequins”). Samples were deeply sequenced on both Illumina short-read and Oxford Nanopore Technologies long-read platforms. Alongside the ground-truth available via the sequins, we created *in silico* mixture samples to allow performance assessment in the absence of true positives or true negatives. Our results show that, *StringTie2* and *bambu* outperformed other tools from the 6 isoform detection tools tested, *DESeq2, edgeR* and *limma-voom* were best amongst the 5 differential transcript expression tools tested and there was no clear front-runner for performing differential transcript usage analysis between the 5 tools compared, which suggests further methods development is needed for this application.

## Introduction

Long-read sequencing technology is becoming more widely used in the field of transcriptomics research. The ability to sequence full transcripts facilitates the study of alternative splicing by identifying and quantifying novel splicing isoforms[1–4] and detecting differences in isoform usage and/or expression between different conditions[5, 6]. A number of customised algorithms for long-read isoform detection and workflows for differential splicing (DS) analysis have been developed in the past few years[7], however, a lack of benchmarking datasets with built-in ground-truth has limited efforts to systematically evaluate the performance of different methods.

Benchmarking studies can help researchers choose the most suitable experimental technologies and computational methods to achieve their study goals[8]. Several benchmark studies for long-read bulk RNA-seq have focused on the comparison of sequencing protocols by analysing data from matched samples. For instance, Soneson *et al*.[9] and Wongsurawat *et al*.[10] compared Oxford Nanopore Technologies (ONT) direct RNA-seq to direct cDNA-seq and Sessegolo *et al*.[11] compared ONT PCR cDNA-seq to direct RNA-seq. In the recent Singapore Nanopore Expression Project (SG-NEx), Chen *et al*.[12] generated a dataset containing 5 human cancer cell lines sequenced by 4 different protocols: ONT direct RNA-seq, direct cDNA-seq, PCR cDNA-seq and Illumina RNA-seq. Synthetic ‘sequins’[13] that contain alternatively spliced transcripts were spiked into some sequencing runs. By comparing different sequencing protocols, they demonstrated the benefit of long-read RNA-seq in transcript-level analysis.

Other long-read benchmarking studies have focused on comparing the performance of different analysis methods. For instance, Dong *et al*.[14] compared 3 isoform detection tools and 2 differential transcript usage (DTU) tools using a mouse neural stem cell dataset and a synthetic sequins dataset. Both experiments had corresponding Illumina short-read data run on equivalent samples that allowed for comparison with ONT long-read data. Sequencing per sample was in the order of a few million reads per sample, which is much lower that the typical number of reads obtained in a short-read RNA-seq dataset. The Long-read RNA-Seq Genome Annotation Assessment Project (LRGASP) Consortium[15] profiled biological triplicates of different human and mouse cell lines (pure and mixed) with human gene derived Spike-In RNA Variants (SIRVs)[16] using different long- and short-read RNA-seq library preparation and sequencing protocols. Human and mouse simulated datasets, and a manatee whole blood transcriptome were also generated. In this study (manuscript in preparation), isoform detection and quantification methods will be compared using the results submitted by challengers. Until now, the systematic performance assessment of differential transcript expression (DTE) and DTU workflows on long-read RNA-seq data has been hampered by small library sizes and the lack of available ground-truth in previous benchmarking studies.

To advance research in this area, we designed and generated a dataset that consisted of matched samples across long-(ONT) and short-read (Illumina) technologies. Each dataset contains 3 replicates from 2 human lung adenocarcinoma cell lines that included sequins spike-in controls. To create biological heterogeneity with inbuilt truth, replicate *in silico* mixtures at 3 different proportions (75:25, 50:50, 25:75) were generated by sampling reads from the different cell line samples. Using these data, we evaluated 6 methods for long-read isoform detection, 5 for DTE analysis and 5 for DTU analysis to guide method selection.

## Results

### Experimental design, benchmarking analysis plan and data quality assessment

Our benchmarking experimental design involved RNA from two distinct human lung adenocarcinoma cell lines, sequins spike-in controls and *in-silico* generated mixture samples (Figure 1a). The cell lines NCI-H1975 and HCC827 were cultured separately in 3 replicates to introduce inter-sample variation due to differences in lab processing and sample preparation to simulate biological variability[17]. Synthetic RNA spike-ins (sequins) from two mixes (A or B) were added to each pure RNA sample to provide inbuilt ground-truth in our experiment. Each pure RNA sample was polyA-selected and cDNA was prepared for sequencing. We used the ONT PCR-cDNA method for long-read sequencing on the ONT PromethION and the NEB Ultra II Directional RNA library preparation for short-read sequencing on the Illumina NextSeq (75bp paired-end), providing data on matched samples across technologies. Reads from each pure RNA sample of a given cell line (denoted as 100:0 and 0:100) were downsampled to a fixed number of reads and combined with the downsampled reads from a corresponding sample of the other cell line in three different proportions (75:25, 50:50 and 25:75) to generate *in silico* mixture samples.

**Figure 1.**
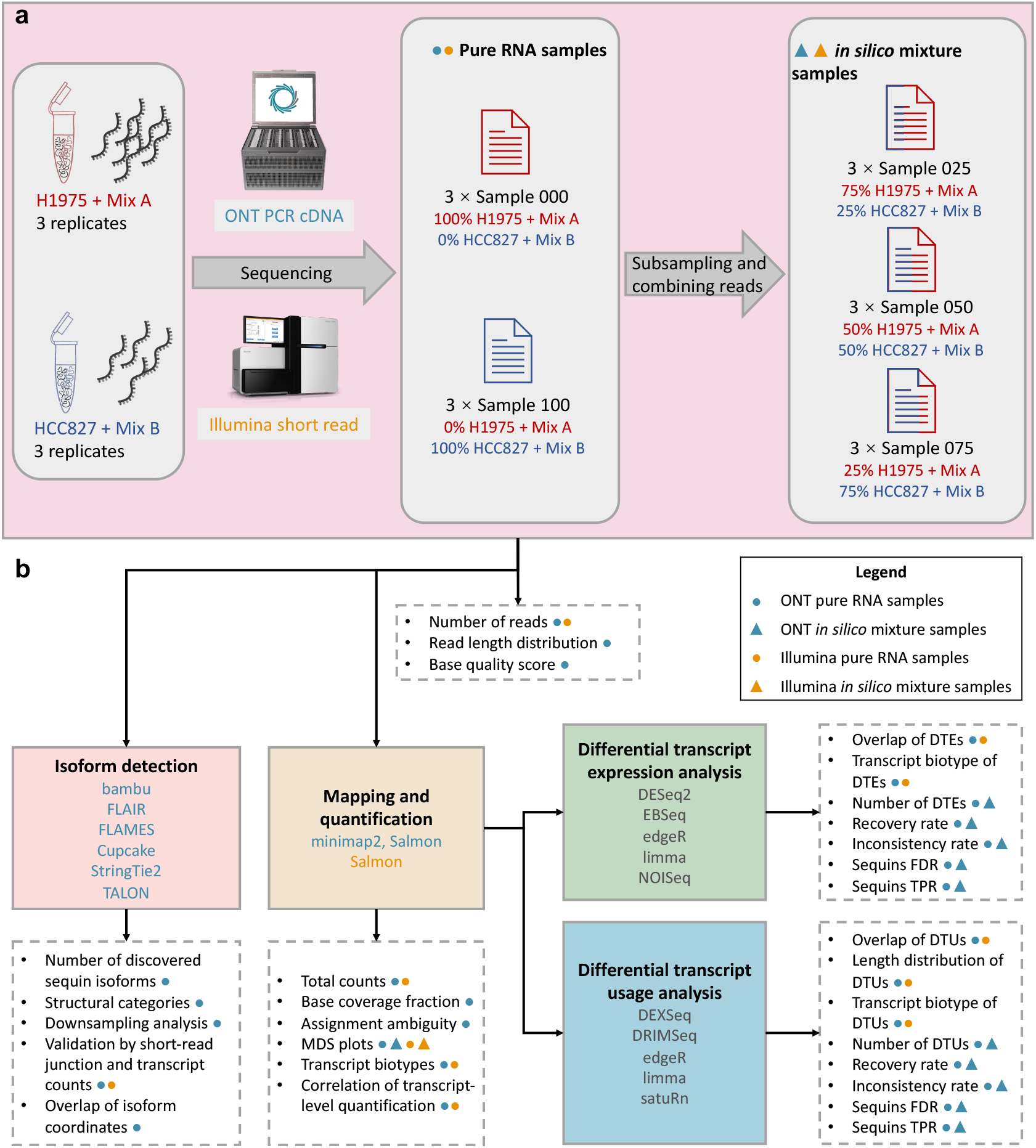
Overview of the experimental design and benchmark analysis. (**a**) Summary of the experimental design involving pure RNA samples obtained from 2 cancer cell line (H1975 and HCC827) with 2 synthetic sequins spike-in control mixes (A and B) and *in silico* generated mixture samples. (**b**) Overview of the analysis workflow to benchmark the performance of different RNA-seq analysis tools for isoform detection and quantification, differential transcript expression (DTE) and differential transcript usage (DTU). Analysis steps and selected methods are shown in shaded boxes with solid borders, while evaluation metrics are listed in boxes with dashed borders.

To validate whether our *in silico* mixture strategy can simulate the variance structure seen in an RNA-seq experiment where physical RNA mixtures were generated, RNA from NCI-H1975+sequins mix A replicate 1 and HCC827+sequins mix B replicate 1, and their *in vitro* mixtures in the same proportions (75:25, 50:50 and 25:75) were sequenced using ONT technology. Additionally, we reanalysed the Illumina short-read data from Holik *et al*.[17] (GSE64098). Triplicates of pure RNA samples of the same cell lines NCI-H1975 and HCC827 as well as lab-generated RNA mixtures in the same proportions (75:25, 50:50 and 25:75) were available. For both datasets, *in silico* mixtures created using the reads from the pure cell line samples were processed together with the pure and lab-mixed RNA samples using the same pipeline. The multi-dimensional scaling (MDS) plots of both the ONT and Illumina data (Supplementary Figure S1a and b) and the high correlation between the counts from lab-mixed and *in silico* mixed samples (Spearman correlation *>*0.93, Supplementary Figure S1)c-e show that the *in silico* mixture samples had very similar gene expression profiles to the lab-mixed samples with the same mixture proportions. This indicates that our *in silico* mixture strategy should be effective at simulating the process of physically mixing RNA in the lab, and saved us 60% of the sequencing costs (only 6 samples needed to be sequenced instead of 15).

By having both ONT long reads and Illumina short reads from the same samples, our experiment provides a useful resource for direct comparison of the characteristics of the different sequencing technologies. Also, the more accurate short reads can be used to verify the isoforms detected by error-prone long-read data. Discrepancies between the results of long- and short-read data analysis can also be examined.

A summary of the analysis workflow is shown in Figure 1b. ONT long reads from pure RNA samples were used for isoform detection using six methods (*bambu, Cupcake, FLAIR, FLAMES, StringTie2* and *TALON*). Both pure RNA samples and *in silico* mixture samples were mapped and quantified against the GENCODE human annotation (Release 33) and sequins annotation (version 2.4). The transcript-level count matrix was used as input to downstream steps such as DTE (five methods: *DESeq2, EBSeq, edgeR, limma, NOISeq*) and DTU (five methods: *DEXSeq, DRIMSeq, edgeR, limma* and *satuRn*). Supplementary Figure S2 shows examples of different DTE and DTU states for 4 genes with 2 transcripts each.

In our ONT dataset, a total of ∼268 million (M) reads were processed and passed quality filtering. The number of reads sequenced from each sample in the ONT dataset varied between 39 M to 51 M (Fig 2a). In comparison, ∼289 M reads were processed in the Illumina dataset. One of the HCC827 samples only had ∼8.6 M reads and was re-sequenced, which delivered a further ∼125 M reads. Overall our experiment contained a total of ∼238 M ONT transcript counts and ∼267 M Illumina transcript counts.

**Figure 2.**
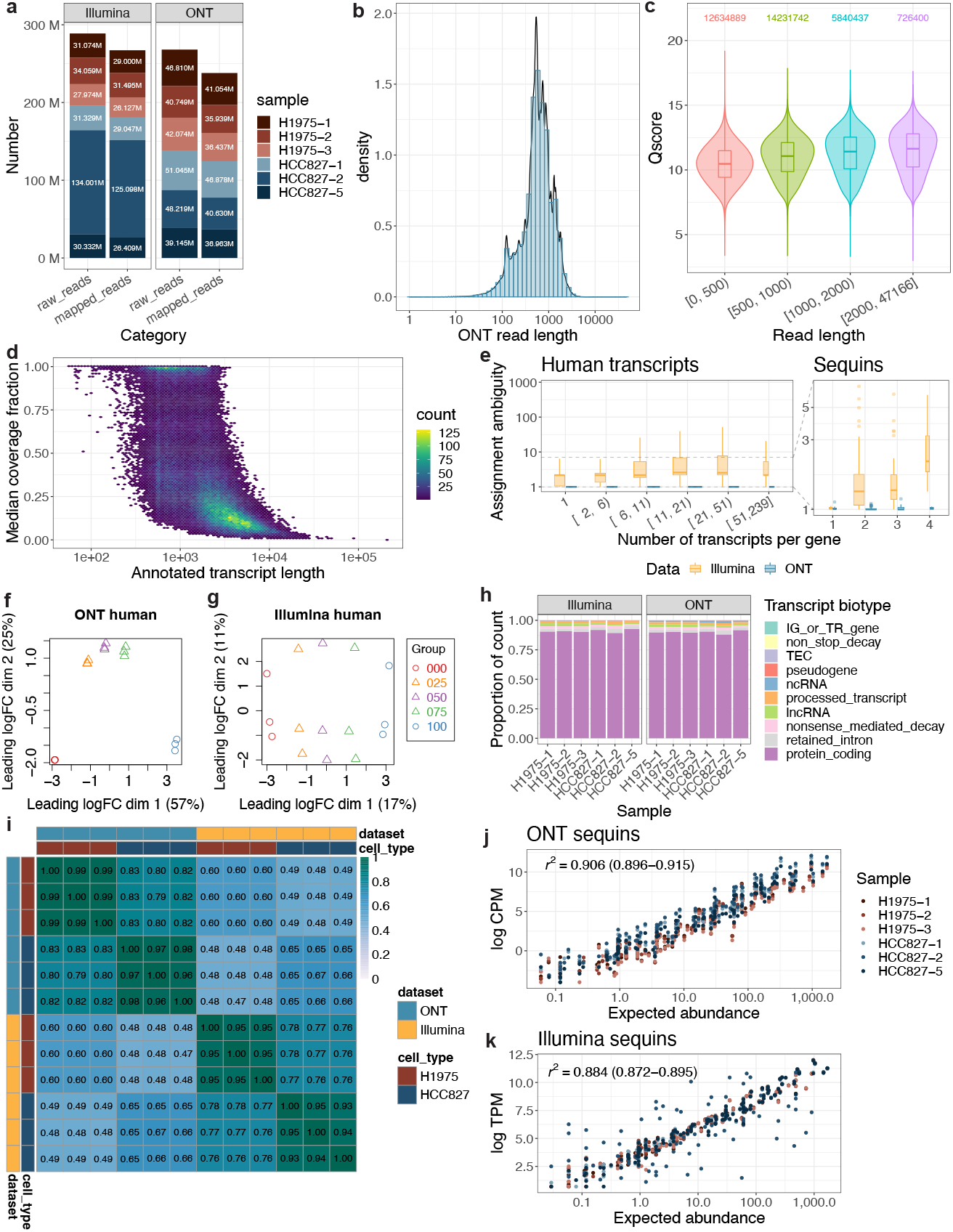
Overview of data quality for the benchmarking dataset. (**a**) The total number of trimmed and demultiplexed reads and transcript-level counts in the ONT and Illumina datasets. (**b**) Distribution of demultiplexed read lengths in the ONT dataset. (**c**) Distribution of read quality in the ONT dataset, splitting by read length. Read quality is defined by the average base quality score of a read. (**d**) A hexagonal 2D density plot showing the correlation between median base coverage rate of a transcript and the length of the annotated transcript. (**e**) Distribution of over-dispersion coefficients in ONT and Illumina dataset from human (left) and sequins (right) transcripts, stratified by number of transcripts per gene. The width of boxes reflect the relative quantity of transcripts within each category. (**f** and **g**) MDS plot showing the pure RNA samples and *in-silico* mixture samples based on human transcript-level logCPM from ONT (**f**) and Illumina (**g**) dataset. (**h**) The proportion of short and long reads assigned to different classes of transcripts in each sample. Transcript-level counts were broken down according to GENCODE’s transcript biotypes annotation. (**i**) Heatmap showing the Spearman correlation of transcript-level CPM (ONT) and TPM (Illumina) from protein coding transcripts and lncRNAs across pure RNA samples. (**j** and **k**) The correlation between observed transcript counts and expected transcript abundance of each sequins transcript from each sample in ONT (**j**) and Illumina (**k**) dataset. The 95% confidence interval for R-squared are shown in parentheses.

In the ONT dataset, median read length is 591 bases, and mean read length is 716.5 bases (Figure 2b), which is similar to Chen *et al*.[12]’s ONT PCR cDNA dataset. Longer reads tend to have slightly higher read quality (Figure 2c). The mean average base quality score for demultiplexed reads is 10.848, which corresponds to an estimated basecalling error rate of 8.2%. The updated ONT basecalling algorithm offers improved accuracy compared to our previous study[14] (estimated basecalling error rate 15.8%).

In short-read RNA-seq, fragmentation during library preparation and read lengths limited to a few hundred bases make it challenging to uniquely map short reads to transcripts, especially for reads from overlapping exons shared by different isoforms of the same gene. In this respect, long-read RNA-seq has a natural advantage since individual reads cover a longer proportion of the transcript. However, current long-read RNA-seq protocols suffer from biases and not all reads correspond to full-length transcripts. Here we refer to the 3’ bias of ONT direct RNA, direct cDNA, PCR cDNA, and PacBio Iso-seq protocols, and other artefacts that can result in truncated reads. A major factor is an incomplete reverse transcription. Exceptionally long or structured RNAs defy the processivity of current reverse transcriptases, and are thus rarely sequenced in full, even in direct RNA sequencing. Size selection (explicitly as part of the standard Iso-seq workflow, or implicitly during the wash steps of ONT protocols) can bias against both short and long transcripts[1, 18]. PCR amplification may also be less efficient for longer transcripts, affecting annotation and quantification, and can introduce GC bias. Finally, library preparation artefacts such as priming on internal polyA stretches, priming on first strand cDNA’s internal polyA stretches, TSO artefacts and concatemers can also lead to reads that do not represent complete transcripts[11].

Here, we calculated the coverage fraction of transcripts by individual ONT reads and Illumina read pairs after mapping them to the reference transcriptome (Supplementary Figure S3). The mean transcript base coverage rate of all ONT reads is 50.4%. As shown in Figure 2d, longer transcripts have lower base coverage rate. Although only 25.7% reads were found to be full length (covering at least 95% of the bases of the corresponding transcripts), 60.3% of transcripts that passed expression filtering have at least one full length read assigned to them, and the longest one among these transcripts is 8,481 base pairs long. By contrast, the mean transcript base coverage rate of all Illumina read pairs is 9.9%. We observed 3’ bias in both our ONT and Illumina dataset such that the 3’ end of the gene body had higher read coverage than the 5’ end (Supplementary Figure S4). The ONT data showed greater decay in coverage over the gene body, but preserved higher coverage at the 3’ end (Supplementary Figure S4).

We estimated the transcript assignment ambiguity for the long and short reads using the over-dispersion calculated by *edgeR*[19]’s *catchSalmon* function. Transcript-specific over-dispersions measure the extra technical variability introduced by the mapping ambiguity during the assignment of reads to transcripts. Transcripts with high sequence homology with other transcripts have greater mapping uncertainty, hence greater over-dispersion coefficients. Over-dispersion coefficients reflect the uncertainty associated with the estimated expression of each transcript, uncertainty that here we compare between short- and long-reads. The over-dispersion for all transcripts are between 1 and 1.15 in our ONT long-read dataset, while two-thirds of the transcripts in the Illumina short-read dataset had an over-dispersion larger than 3.56 (Figure 2e). Importantly, transcripts from genes with more isoforms tend to have significantly higher over-dispersion in the Illumina dataset. Due to the lower complexity of the sequins transcriptome, the mean over-dispersion of sequins (1.688) was lower than human transcripts (6.184). This suggests that the ONT PCR cDNA sequencing protocol far outperforms short-read sequencing in assigning reads to specific transcripts, especially between multiple isoforms of the same gene, despite current limitations in obtaining full-length reads.

To examine the variability in the data, MDS plots were generated after low-expression filtering, normalization and converting the read counts from all samples to log_2_CPM (counts per million). Samples were clearly separated on dimension 1 by mixture proportion in both the ONT and Illumina datasets (Figure 2f and g). Samples from the same mixture proportion also appeared closely on the MDS plot of human transcripts in ONT dataset (Figure 2f). In the MDS plots for sequins transcripts for both datasets (Supplementry Figure S5a and b), dimension 1 explained most variance in the data (80% for ONT, 79% for Illumina). Overall, the MDS plots show that the major variance in the data can be explained by the biological difference between different mixture proportions of different cell lines. We also calculated the biological coefficient of variation (BCV) for transcripts on both datasets. As shown in Supplementary Figure S6, the biological variation level as suggested by common BCV of human transcripts on the Illumina short-read dataset (0.54) was higher than the ONT long-read dataset (0.20), while the BCVs of all sequin transcripts were low on both datasets (tagwise BCVs 0.0023 to 0.0067 for ONT, 0.0006 to 0.0049 for Illumina).

In order to compare the read distribution across different RNA classes from long- and short-read sequencing, we annotated the count matrices using the transcript biotype information available from the GENCODE annotation. The proportion of reads assigned to most biotypes in each pure RNA sample (Figure 2h) were very similar between ONT and Illumina datasets; about 90% of reads were assigned to protein coding transcripts in both datasets. Our ONT long-read sequencing captured marginally more non-coding RNAs (ncRNAs). To compare the transcript-level quantification obtained from ONT long-read and Illumina short-read sequencing, we calculated the Spearman correlation of long-read CPM and short-read TPM across each pure RNA sample (Figure 2i, Supplementary Figure S7). We noticed that some transcripts which were detected to be highly expressed in one platform, only had limited number of reads detected in another platform (Supplementary Figure S7a). The transcript-level quantification from protein coding transcript and lncRNAs showed very high correlation (*>*0.93) between replicates. The Spearman correlations of quantification from the same cell lines sequenced on different platforms (0.60 to 0.67) are lower than that of different cell lines on the same platform (0.76 to 0.83), indicating the existence of strong batch effects between different sequencing platforms (Figure 2i). The inconsistency of quantification between platforms is stronger for transcripts in other biotypes (Spearman correlation 0.28 to 0.35, Supplementary Figure S7b), which leads to reduced cross-platform correlation when calculated using all transcripts (Supplementary Figure S7c). To investigate whether the different correlations observed by biotypes are related to the difference in their expression levels, we stratified the transcripts into three expression level groups by the 1/3 and 2/3 quantiles of their total counts (i.e. low - bottom 1/3, medium - middle 1/3, high - top 1/3) and calculated the composition of biotypes and the between-sample Spearman correlations in each group. The proportion of protein coding transcripts were higher in the medium- and high-expressed groups (Supplementary Figure S8a). Interestingly, the correlations of quantification from high- and medium-expressed transcripts between samples from the same dataset (Supplementary Figure S8b and c) were similar to the correlations calculated using all transcripts (Supplementary Figure S7d), whether between replicates (*>*0.90) or between samples from different cell lines (0.75 to 0.84). However, the correlations of quantification between samples sequenced on different platforms were low across all expression level groups (0.12 to 0.29 for highly expressed transcripts, 0.14 to 0.32 for medium expressed transcripts, and -0.44 to -0.16 for lowly expressed transcripts, Supplementary Figure S8b-d). By contrast, the correlation of sequins transcript quantification across sequencing platforms is very high (0.96 to 0.97, Supplementary Figure S7d).

To evaluate the accuracy of quantification, we compared our transcript-level quantification of the sequins spike-in transcripts to their annotated abundance. Both CPM (effective TPM) from ONT long reads (Figure 2j) and TPM from Illumina short reads (Figure 2k) had strong positive linear relationships with the annotated abundance. The linear relationship for the ONT data is slightly stronger than that of Illumina data, indicating that improved transcript-level quantification can be obtained using long-read sequencing.

### Comparisons of isoform detection methods

We tested six isoform detection and quantification methods for long-read data: *bambu*[20] (https://github.com/GoekeLab/bambu), *Cupcake* (https://github.com/Magdoll/cDNA_Cupcake), *FLAIR*[21], *FLAMES*[22], *StringTie2*[23] and *TALON*[24]. Using the sequins, we evaluated how well the methods recovered known transcripts in the original full dataset using the complete annotation, and how well they discovered known, but missing (‘novel’) transcripts in a downsampled dataset using an annotation where selected isoforms (40 in total) were removed at random (see Methods). Ideally, all known sequins transcripts should be detected with no novel isoforms. Our results show that *bambu* generated the highest recall (∼0.91) and precision (∼0.62) rates, detecting the most sequins transcripts and the fewest artefactual isoforms in the full dataset (Figure 3a). *StringTie2* performed the best in both such aspects in the downsampled analysis, recovering (80%) of the removed, and (∼76%) of the retained isoforms with the highest precision. *Cupcake, FLAIR* and *TALON* had the lowest precision rates across all analyses, having found a significantly higher number of artefactual isoforms than other methods. *FLAMES* had a similarly low recall rate for the detection of isoforms in the downsampled analysis to *FLAIR* and *TALON*, particularly of the removed isoforms (Figure 3b, bottom right panel). Interestingly, however, it had the second highest precision rates after *StringTie2*, for both retained and removed sequin isoforms, indicating relatively high accuracy to detect true isoforms. The tools with the highest recall rates for sequin recovery and the only ones to recover at least half of the removed sequins were *bambu* (0.65), *Cupcake* (0.80) and *StringTie2* (0.80); the worst-performing tool in this regard was *FLAIR* (∼0.28) (Figure 3b, bottom right panel). However, *FLAIR* produced the highest recall rate (~0.63) after *StringTie2* (∼0.76) and *bambu* (∼0.72) for retained isoforms. Overall, *bambu* and *StringTie2* outperformed all other methods in the full and downsampled sequins analyses, respectively.

**Figure 3.**
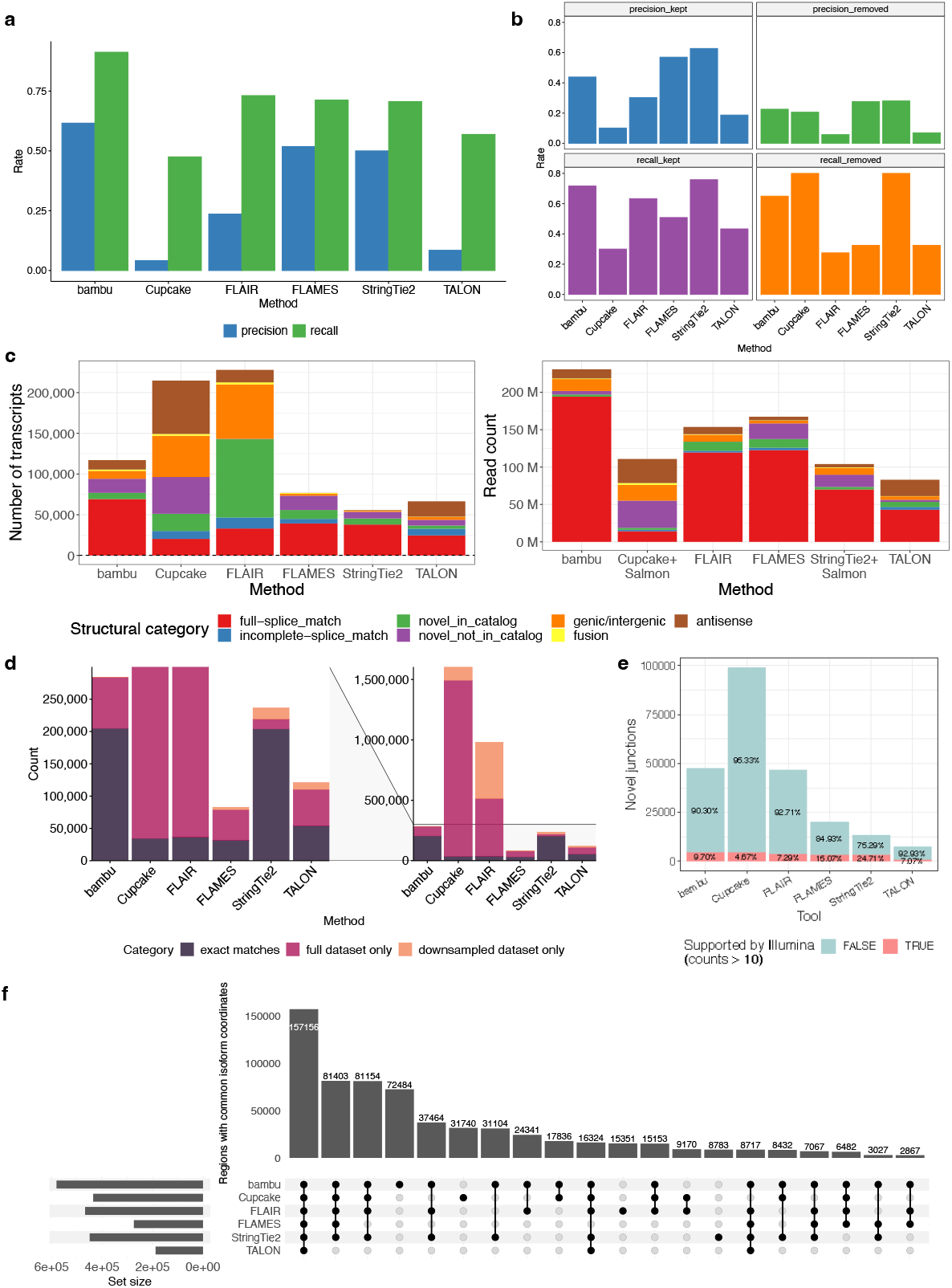
Comparison of isoform identification and quantification methods using pure RNA samples. (**a**) Precision and recall rates of sequins in the reference annotation detected by each tool in the full ONT data. (**b**) Precision and recall rates of sequins that were *kept* (left-hand panel) or *removed* (right-hand panel) from the reference annotation by each tool in the downsampled ONT data. **(c)** Bar plots showing the number of isoforms detected in the ONT data (left) and the number of counts from isoforms (right), coloured by isoform structural categories. Known transcripts (i.e. true positive) constitute the red “full splice match” category; other categories represent novel transcripts detected. **(d)** A bar plot showing the the number of isoforms detected in both the full and downsampled ONT long-read datasets (“exact matches”), in the full dataset only and in the downsampled dataset only. (**e**) Percentage of the novel junctions from isoforms detected by each method that were supported by more than 10 Illumina counts for each tool. Redundant junctions were excluded. (**f**) An UpSet plot showing the 20 largest intersections of isoform coordinates between isoform annotations identified by each tool in the ONT long-read data.

Exploration of the transcripts detected in the human cell line RNA revealed disproportionately large numbers of novel transcripts detected by *FLAIR* and *Cupcake* as indicated by the number of transcripts belonging to isoform structural categories other than “full splice match” (Figure 3c). The highest percentage of known transcripts, represented by the “full splice match” category, was produced by *StringTie2* (∼68%). For reads assigned to transcripts in each structural category, most counts were from known isoforms in all tools, except *Cupcake*. Overall, *bambu* detected the largest number of known transcripts (69,252) and yielded the highest read counts for known transcripts (∼194 M). In contrast, *Cupcake* generated the smallest number of known transcripts (20,334), with the lowest read counts for this category (∼14 M), and the highest proportions of novel transcripts (∼91%) and read counts (∼87%). Of note, *bambu* retained all *a priori* annotated isoforms in the output GTF annotation file even if some of them did not meet the criteria to be identified as ‘detected’ in the dataset. Its subsequent use of all of pre-existing isoforms for quantification makes annotated isoforms easier to be considered as ‘detected’ in our analysis.

We also compared isoforms identified in the full (100%) versus the downsampled (10%) cell line datasets, to evaluate the performance of isoform detection at lower sequencing levels (Figure 3d). Ideally, as many isoforms as possible detected in the full dataset should be recovered completely as exact matches to the annotations generated from the downsampled dataset, with no novel isoforms in the downsampled dataset that were not detected in the full dataset. *StringTie2* and *bambu* showed superior performance compared to other methods and were the only methods to yield exact matches for more than 70% of the isoforms detected in both datasets. *StringTie2*, with 236,428 total comparisons, recovered the highest proportion of exact matches (203,776, ∼86%), making it the best-performing tool in this analysis. It also produced the lowest proportion of isoforms detected only in the full dataset (15,482, ∼7%). *bambu* recovered ∼72% (204,633) of exact matches and produced the lowest proportion of isoforms detected only in the downsampled dataset (∼0.03%), with just 78 isoforms. In contrast, results from the worst-performing tools in this analysis, *FLAIR* and *Cupcake*, comprised proportions of exact matches of less than 5% (36,528/980,929 and 34,362/1,604,301, respectively). *FLAIR* also identified 466,783 isoforms in the downsampled dataset only, the highest proportion (∼48%) out of all tools. A large number of isoforms detected by *Cupcake* (1,458,931 or ∼91%) were unique to the full dataset.

In an attempt to distinguish between true positive transcripts and false positives, we examined the level of support provided by high accuracy short reads[25] by looking at the percentage of novel junctions from isoforms detected by each tool that were supported by Illumina reads (*>*10 uniquely mapped junction counts, Figure 3e). *StringTie2* performed the best in this analysis and was the only method that have more than 20% of novel junctions supported by short reads. *FLAMES* had the second highest proportion of supported junctions. *FLAIR* and *bambu* detected similar high number of unsupported novel junctions and low proportion of supported novel junctions. *Cupcake* detected the highest number of novel junctions, but 95.33% of them were not supported by Illumina reads. Both the number of novel junctions supported or not supported by Illumina reads detected by *TALON* were the lowest among all tools we compared. We also quantified the transcripts detected by each tool using Illumina short reads (Supplementary Figure S9). Theoretically, there should be a high degree of agreement between the long and short reads among isoforms; therefore false junctions or transcripts attributable to long-read sequencing error should not be assigned supporting short reads. The correlation for *StringTie2* between ONT long-read log_2_CPM (counts per million) and Illumina short-read log_2_TPM (transcripts per million) was also highest (R = 0.58). *FLAIR* and *bambu* yielded similar correlations (*R* = 0.54 and *R* = 0.53, respectively), but detected many more transcripts (n = 228,204 and n = 151,126, respectively) than *StringTie2* (n = 55,630). The correlation for *FLAMES* was close to 0.50 (*R* = 0.47). The correlations of quantification between ONT and Illumina data for *TALON* and *Cupcake* were significantly lower (*R* = 0.20 and *R* = 0.23, respectively).

To examine the intersection of results from all isoform methods at the genome level, we developed a genome binning approach. Due to differences in transcript naming conventions and output formats, the coordinates of the isoforms detected by each tool were used instead. We compared the overlap of coordinates in two kilobase bins along the genome. In the UpSet plot shown in Figure 3f, the largest intersection of bins with common isoform coordinates encompasses all tools, indicating a high degree of agreement among the methods for isoforms detected. The next two largest intersections also include most tools; the second largest intersection excludes *TALON* and the third excludes *FLAMES* and *TALON*. The fourth intersection shows that many unique regions overlap with the coordinates of a subset of isoforms detected only by *bambu*, as does the sixth intersection with *Cupcake*, suggesting that a sizeable number of unique isoforms were detected by these tools.

### Comparisons of differential transcript expression methods

Given the advantages of long-read sequencing in terms of lower transcript assignment uncertainty and better transcript-level quantification, traditional differential gene expression analysis can be extended to the transcript-level by applying differential expression (DE) methods to transcript count matrices[26, 27]. We evaluated five popular DE analysis methods including *DESeq2* [28], *edgeR*’s quasi-likelihood pipeline[19], *EBSeq* [29], the *limma-voom* pipeline[30] and *NOISeq* [31], using the transcript count matrix as input.

Here, we applied the same DTE analysis workflows on transcript counts from both ONT long-read and Illumina short-read data, and compared the differentially expressed transcripts detected between pure RNA samples (100 vs 000) by different methods on each dataset. For transcripts tested across both the Illumina and ONT datasets, results from different methods showed high concordance, with the largest intersection of differentially expressed transcripts detected by all methods across both datasets (Figure 4a, Supplementary Figure S10a). The *t* -statistics calculated by *limma-voom* pipeline and converted to *Z*-scores had high correlation between the ONT and Illumina data (Figure 4b). The length distribution of detected DTEs were similar between the methods across both datasets (Supplementary Figure S10b). If we consider all transcripts tested by any method in either the Illumina or ONT datasets (variation here arises due to differences in filtering), we observe many that were specifically detected as differentially expressed in the ONT dataset by one or more methods (Supplementary Figure S10a), and that all methods reported more DTEs in the ONT dataset than in the Illumina dataset (Supplementary Figure S10a). In both datasets, *NOISeq* detected more than 3,000 unique DTEs which were not detected by other methods in any dataset. More lncRNAs and retained introns were detected to be significantly differentially expressed in the ONT dataset compared to the Illumina dataset (Figure 4c, Supplementary Figure S10c).

**Figure 4.**
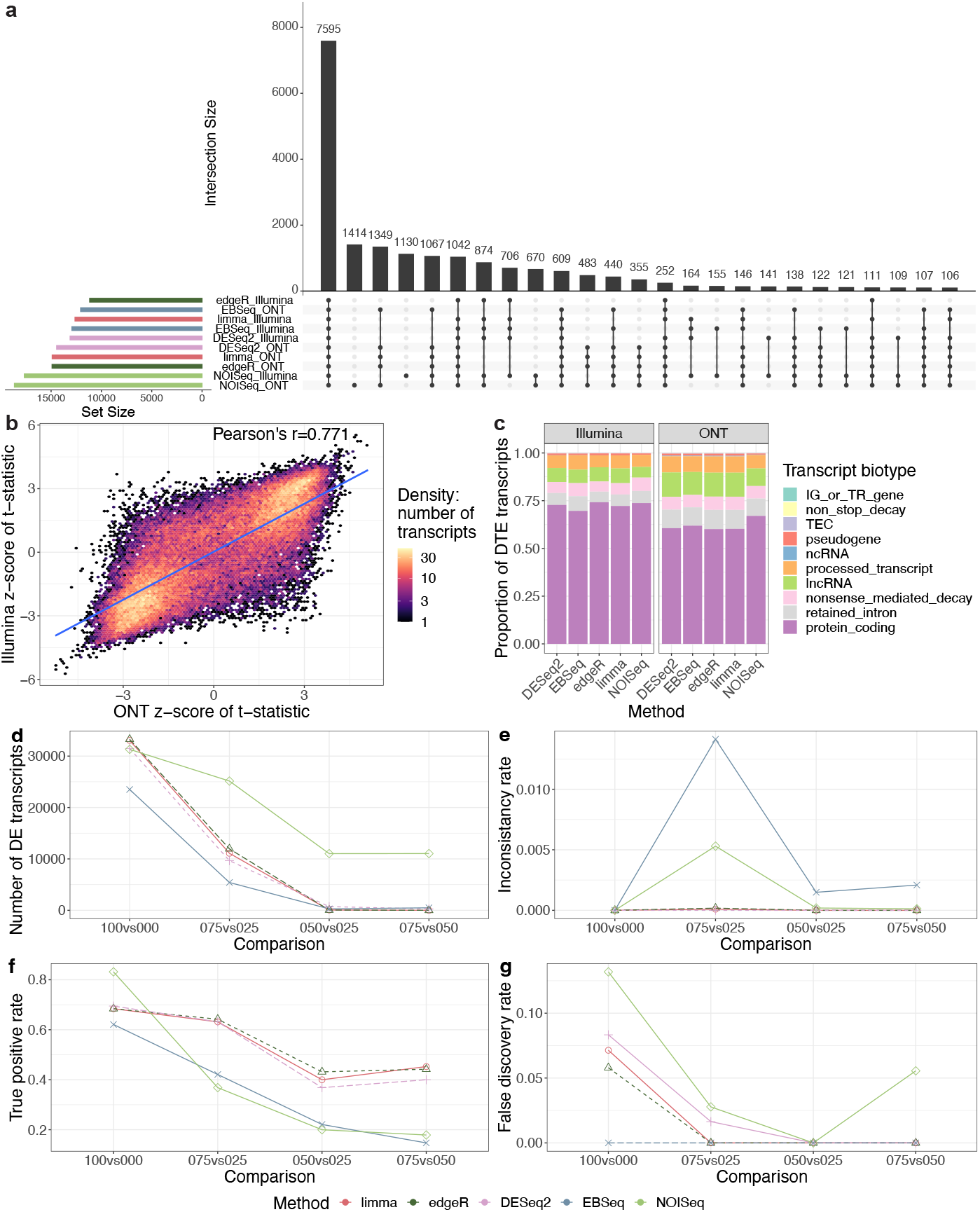
Comparisons of differential transcript expression methods using *in silico* mixtures. (**a**) An UpSet plot showing the 25 largest intersections of differentially expressed transcripts between HCC827 (100) and H1975 (000) samples detected by each tool in Illumina short-read and ONT long-read data. Only transcripts tested by all methods on both Illumina and ONT datasets are included. (**b**) A hexagonal 2D density plot of the *Z*-score standardized *t* -statistics calculated between HCC827 (100) and H1975 (000) samples from Illumina short-read and ONT long-read data. (**c**) The proportion of differentially expressed human transcript biotypes detected by each tool in Illumina short-read and ONT long-read data. (**d**) The number of differentially expressed human transcripts detected in each comparison of the ONT data showing the power and the ability to recover discoveries from the 100 versus 000 comparison of each method. (**e**) The rate of differentially expressed human transcripts detected that were inconsistent with discoveries from the 100 versus 000 comparison of each method. (**f**) The true positive rate (TPR) of different comparisons from each differential transcript expression tool in the ONT sequins data. (**g**) The false discovery rate (FDR) of different comparisons from each differential transcript expression tool in the ONT sequins data.

To further evaluate the performance of DTE methods, various metrics relying on the *in silico* mixture series were calculated. Relative to the comparison between pure RNA samples (100 vs 000), the differences between the comparison 075 vs 025 samples is less pronounced, while the comparisons 050 vs 025 and 075 vs 050 will be more subtle again due to the more similar mixing proportions. Although the true positive or true negative DTE transcripts between the cell lines are not annotated, based on the fact that the same set of transcripts must be differentially expressed between any pair of samples in a mixture design, we were able to estimate the sensitivity and precision of methods using the following metrics: the number of DTEs detected in each comparisons, recovery rate (proportion of DTEs detected in 100 vs 000 that are also detected in other comparison), and inconsistency rate (proportion of DTEs in comparisons of mixture samples that are not detected in the 100 vs 000 comparison). Furthermore, the known differences in transcript abundance between sequins mixtures A and B allowed us to calculate the false discovery rate (FDR) and true positive rate (TPR) of DTE methods.

Using an adjusted *p*-value cutoff of 0.05 (or estimated probability of DE *>*0.95), *limma* and *edgeR* detected the largest number of DTEs (∼33,000) between pure RNA samples (100 vs 000) among a total of ∼60,000 human transcripts, followed by *DESeq2* and *NOISeq* reporting a bit less DTEs (∼31,000, Figure 4d). Among the five methods we compared, *NOISeq* achieved the highest recovery rate for human genes across the various comparisons of mixture samples (Supplementary Figure S10d). *DESeq2, edgeR* and *limma* performed similarly in terms of the total number of DTEs they obtained (Figure 4d) and their recovery of DTEs found in 100 vs 000 comparison in the other mixture comparisons (Supplementary Figure S10d). *EBSeq* detected the least number of DTEs from the 100 vs 000 comparison, and it also had the lowest recovery rate in the 075 vs 025 comparison (Figure 4d, Supplementary Figure S10d). The inconsistency rate of all methods are below the 0.05 threshold (Figure 4e), with *limma, edgeR* and *DESeq2* being very close to zero, indicating that all methods controlled false discoveries well in the human ONT data.

For the sequins data, *NOISeq* detected the most total and true positive DTEs in the 100 vs 000 comparison (Supplementary Figure S10e, Figure 4f), but it had the lowest recovery rate for all other mixture comparisons (Supplementary Figure S10f), and the highest FDR among all methods (Figure 4g). *DESeq2, edgeR* and *limma* outperformed the other tools in terms of recovery rate (Supplementary Figure S10e), however, their FDR in the 100 vs 000 comparison exceeded the nominal 0.05 threshold (Figure 4g), which indicates the potential for more false discoveries than expected. Although *EBSeq* had a relatively lower power to detect DTEs (Figure 4d), it was the only tool that controlled FDR well in all comparisons (Figure 4g). Similar results were observed on the Illumina dataset, but *edgeR* and *DESeq2* had no FDR exceeding the 0.05 threshold in any comparison (Supplementary Figure S11).

Library size is a key factor that can influence the results of a DE analysis, with more reads per sample generally increasing power to detect differentially expressed genes or transcripts, albeit at higher cost per sample. To explore how DTE results change with varying read numbers per sample and how many long reads per sample were required to provide comparable DTE results to short-read studies, we downsampled each pure long- and short-read RNA sample to obtain new datasets in which each sample had a fixed read number (1, 3, 5, 10, 15 or 20 million). DTE analyses were performed on each dataset using *edgeR*, and the results were compared to the full datasets that had not undergone downsampling. For both Illumina and ONT, larger fold-changes between pure NCI-H1975 and HCC827 samples were observed from datasets with larger library sizes in dimension 1 of the MDS plots (Supplementary Figure S12a-b). More differentially expressed transcripts were detected in datasets with more reads, while more DTE was always detected on the ONT datasets compared to the Illumina datasets with the same library sizes (Supplementary Figure S12c-d). On both ONT and Illumina datasets, *edgeR* performed similarly in terms of its DTE recovery rate calculated using results from the full dataset as the reference (Supplementary Figure S12e). Interestingly, we found that 10 million reads per sample was sufficient to recover more than half of the differentially expressed transcripts detected in the full datasets, and the ONT data with this number of reads had the power to detect similar numbers of differentially expressed transcripts compared to the full Illumina dataset (Supplementary Figure S12c-e). Although the total numbers and proportions of inconsistent differentially expressed transcripts in the downsampled ONT datasets were higher than the equivalent Illumina datasets across different library sizes, the number of inconsistent discoveries amongst the top ranked transcripts was consistently lower across all ONT datasets, especially those with larger library sizes (Supplementary Figure S12c, f-g).

### Comparisons of differential transcript usage analysis methods

Other than studying the difference of transcript expression levels between experimental groups, it is also of biological interest to study differential splicing (DS). One way to study DS is through differential transcript usage (DTU) analysis, where the relative changes in proportion of expressed transcripts from a given gene are examined between conditions. Differential exon or exon junction usage (DEU) analysis is an alternative way to study DS that focuses on the usage changes for a given gene at the exon-level. If transcript-level counts rather than exon-level counts were used as input, models designed for DEU analysis can also be used to perform DTU analysis[14, 32]. We selected five methods to perform DTU analysis, including two DTU methods (*DRIMSeq* [33], *satuRn*[34]) and three methods that were originally designed for DEU analysis (*DEXSeq* [35], *limma-diffSplice* pipeline[30], *edgeR-diffSliceDGE* pipeline[19]).

To study DTU events, there are two ways to identify significant changes between different experimental groups. The transcript-level test measures whether each individual transcript changes in proportion between conditions, while the gene-level test examines whether, for a given gene there are any transcripts with DTU. The transcript- and gene-level DTU results are usually combined in practice to identify features of biological interest. Here, we compared the DTU transcripts and genes detected between the pure (100 vs 000) samples by different methods on each dataset. All methods except for *limma-diffSplice* had more significant findings in the ONT data than the Illumina data at both the transcript-(Supplementary Figure S13a) and gene-level (S14a). One reason for this is that a lot of significant DTU features detected in the ONT dataset did not pass expression filtering in the Illumina dataset (Figure 5a, Supplementary Figure S14b). Among all the DTU features detected by any method, only 14.4% of transcripts (Supplementary Figure S13b) and 23.1% of genes (Supplementary Figure S14c) were detected to be significant on both the ONT and Illumina dataset. The inconsistency of DTUs detected in the ONT and Illumina datasets was higher at the transcript-rather than the gene-level, both before (Supplementary Figure S13b and S14c) and after (Supplementary Figure S13c and S14d) filtering out transcripts or genes that were only tested in one dataset. All methods tend to find shorter DTU transcripts in the ONT dataset than the Illumina dataset (Figure 5b), while the difference in length of DTU genes detected in ONT and Illumina dataset is subtle (Supplementary Figure S14e). A higher proportion of DTU genes and transcripts that are not of the *protein coding* biotype were detected on the ONT dataset compared to the Illumina dataset, especially for the *lncRNAs* and *processed transcripts* classes for the transcript-level analysis (Figure 5c), and *lncRNAs* for the gene-level analysis (Supplementary Figure S14f).

**Figure 5.**
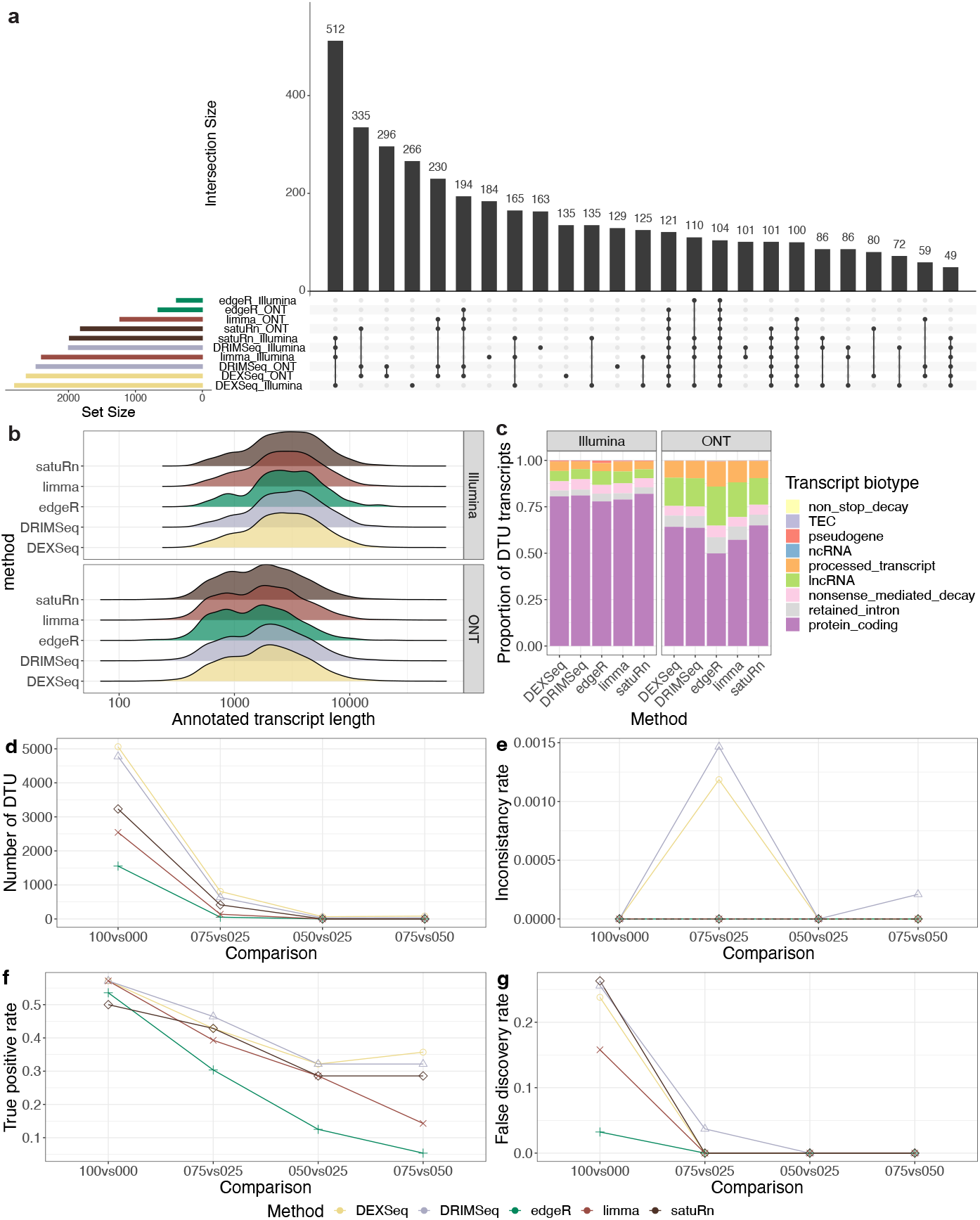
Transcript-level comparisons of differential transcript usage methods using *in silico* mixtures. (**a**) An UpSet plot showing the 25 largest intersections of DTU features between HCC827 (100) and H1975 (000) samples detected by each tool in Illumina short-read and ONT long-read data. Only transcripts tested by all methods on both Illumina and ONT datasets are included. (**b**) Distribution of annotated length of DTU features detected by each method in Illumina (top) and ONT (bottom) data. (**c**) The proportion of DTU features of different biotypes detected by each tool in Illumina short-read and ONT long-read data. (**d**) The number of DTU features in each comparison of ONT data showing the power and the ability to recover discoveries from the 100 versus 000 comparison of each method. (**e**) The rate of detected differential used human transcripts inconsistent with discoveries from 100 versus 000 of each method. (**f**) The true positive rate (TPR) of different comparisons from each DTU analysis tool in the ONT sequins data. (**g**) The false discovery rate (FDR) of different comparisons from each DTU analysis tool in the ONT sequins data.

We evaluated the performance of the five DTU methods in a similar manner to the DTE methods, by calculating the power, recovery rate and inconsistency rate of each method at the transcript- and gene-level using the ONT *in silico* mixture series, and calculating the true positive rate and false discovery rate for the sequins transcripts and genes. *DRIMSeq* and *DEXSeq* detected the largest number of human DTU transcripts (Figure 5d) and genes (Supplementary Figure S15a), while *edgeR* detected the least DTUs. The power of all DTU methods declined dramatically as the mixing proportions compared became more similar, with recovery rates below 0.2 for all methods for both human transcripts (Supplementary Figure S13d) and genes (Supplementary Figure S15b). The recovery rate of *edgeR* and *limma* were slightly lower than other methods. The inconsistency rate of all methods are below the 0.05 threshold (Figure 5e and S15c).

For the ONT sequins dataset, *DRIMSeq* and *limma* detected the largest number of DTU transcripts and genes for the 100 vs 000 comparison (Figure 5f, Supplementary Figure S13e, Supplementary Figure S15d and e), but the recovery rate of *limma* was lower than *DRIMSeq* at both the transcript-(Supplementary Figure S13f) and gene-level (Supplementary Figure S15e). *DEXSeq* also detected the largest number of DTU sequins transcripts, and had the highest recovery rate for the comparison 075 vs 050 (Supplementary Figure S13f). The FDR of *DRIMSeq, satuRn* and *DEXSeq* in the 100 vs 000 comparison were all above 0.2 at both the transcript-(Figure 5g) and gene-level (Supplementary Figure S15g). Although *edgeR* was the only tool that controlled the transcript-level FDR to be lower than the cutoff 0.05 in the 100 vs 000 comparison (Figure 5g), and had the lowest gene-level FDR (Supplementary Figure S15g), its power to detect DTU at the transcript- or gene-level was lowest among all tools for all comparisons between the *in silico* mixture samples (Figure 5f, Supplementary Figure S15d).

## Discussion

By introducing ground-truth in the form of sequin spike-ins, we could assess the performance of tools at each step of the workflow using a small number of genes and transcripts in the presence of technical variation only. Furthermore, the *in silico* mixtures provided a means to broadly assess the performance of DTE and DTU tools on a genome-wide scale, where true positives and negatives are unknown and both technical and biological variation was present. Our *in silico* mixture strategy provides extra levels of ground-truth without extra cost, and can be easily applied to other datasets that have replicate samples across multiple groups.

Previous studies of ONT long-read RNA-seq data contained low read numbers per sample resulting in a loss of power for DE analyses[9, 14, 27]. Compared to other long-read RNA-seq studies such as the SG-NEx project[12] (34 million long reads per cell line on average) and the LRGASP project[15] (3-60 million reads per cell line per sequencing protocol), our samples were sequenced to a greater extent using one single long-read protocol (134 million reads per cell line on average, Figure2a). Our results suggest that the sequencing level per samples we obtained can support the detection of both novel isoforms (Figure 3) and a large number of DTE (Figure 4 or DTU 5). Using a downsampling strategy, we showed that having more reads per sample results in more DTE detected in both the ONT and Illumina datasets, although our ONT long-read dataset provided better power for DTE analysis compared to Illumina short-read data using the same number of reads (Supplementary Figure S12).

The PCR cDNA-seq protocol from ONT yields a greater number of reads than other ONT protocols[11, 12, 15], however, the reads can be truncated due to internal priming in both first-strand and second-strand cDNA synthesis [11]. This results in shorter read lengths than have been observed using either direct RNA-seq and direct cDNA-seq protocols, which also suffer from partial transcript coverage themselves [12]. This limits our ability to obtain reads covering full-length transcripts (Figure 2d, Supplementary Figure S3). Further protocol improvements will be required to ensure reliable information can be obtained for transcripts above 2 kb in length such as those from *XIST* (16.5 kb) and *TITIN* (80 kb). The use of updated basecalling algorithms has seen a reduction in sequencing error rate in these data (Figure 2c) compared to previous studies[14], although it is still considerably higher than the error rate of short-read sequencing.

We were able to obtain reliable transcript quantification on both ONT and Illumina data for sequins transcripts using *Salmon* (Figure 2j and k, Supplementary Figure S7d). However, transcript quantification using short-read data was much more challenging for the human data due to its complexity. The higher transcript assignment ambiguity in the Illumina data (Figure 2e), leads to increased technical variation in the Illumina human transcript counts, as reflected in MDS plots where the proportion of variance explained by dimension 1 that separated samples by groups was lower (17%), and replicates from the same group were not clustered together on dimension 2 (Figure 2g). Although a large proportion of isoforms (72% or more) were detected in common between the full and downsampled dataset by both *StringTie2* and *bambu* (Figure 3d) which suggests the possibility of hybrid analysis using low-depth long-read and high-depth short-read sequencing, it is still challenging to obtain accurate short-read transcript-level quantification with current methods.

Due to the limitations of current long-read RNA-seq technology, it is challenging to distinguish true novel isoforms from false discoveries. If isoform filtering is too stringent, many genuine isoforms will be missed. Both the existence of false discoveries and missing true isoforms can impact the accuracy of transcript-level quantification. Our comparison of isoform detection tools showed that *StringTie2* and *bambu* outperformed other tools in general on ONT data. *StringTie2* showed better accuracy, while *bambu* showed better power to detect novel isoforms and assignment of reads to produce counts (Figure 3). However, even for relatively simple cases like the sequins, the best performing tools still showed mostly low precision rates, especially in the downsampled analysis (∼0.10 to 0.63 for transcripts retained in the annotation, and ∼0.07 to 0.28 for transcripts that were deleted from the annotation). Even for the best performing tools, less than 25% of the novel junctions detected on the ONT dataset were supported by high-accuracy short reads (Figure 3e). This suggests that current long-read isoform detection tools overpredict novel isoforms, which presents an opportunity for methods development to improve this situation. Users of current tools should be mindful of this, and consider orthogonal approaches to validate the authenticity of any novel transcripts uncovered. For ease of comparison, we mostly relied upon the default settings of each method for identifying novel isoforms. However, the large number of artefactual sequin isoforms, novel transcripts and junctions that were not supported by short-read data detected by *Cupcake, FLAIR* and *TALON* (Figure 3) suggests that these parameters may not be optimal for datasets with large numbers of reads.

For long-read RNA-seq, our study is the first to compare DTE and DTU methods on a controlled dataset with a tens of millions of reads per sample, as is typically available in short-read studies. Our work therefore complements the efforts of other long-read RNA-seq benchmarking projects such as SG-NEx[12] and LR-GASP project[15]. *DESeq2, edgeR* and *limma-voom*, which are considered as ‘gold standard’ methods for short-read DE analysis, showed similarly good sensitivity to detect differentially expressed transcripts on our long-read dataset, while *edgeR* outperformed other methods in terms of false discovery control (Figure 4, Supplementary figure S10). However, for DTU analysis, none of the methods we compared had a good balance between power to detect DTUs and control of false discoveries (Figure 5, Supplementary figure S13, S14 and S15), which suggests there is still a gap to be filled to develop DTU analysis tool that are better tailored to long-read RNA-seq data. *DEXSeq* and *DRIMSeq* showed higher sensitivity in most cases, but their false discoveries were not controlled for the sequin spike-ins. In contrast, *edgeR-diffSpliceDGE* performed better in terms of FDR control, but had the lowest power to detect DTU genes and transcripts. Unlike DTE analysis where each transcript is tested independently, DTU analysis calculates the proportion of transcript expression relative to all transcripts from the gene, which can be impacted more readily by changes in quantification of any transcript from a given gene. Therefore, the difference of quantification in ONT and Illumina data (Figure 2i, Supplementary Figure S7) had a larger impact on DTU results (Figure 5a) than on DTE results (Figure 4a and b).

Our analysis has included many popular tools and covered the most popular use cases of long-read RNA-seq. Methods for other analysis tasks such as read alignment, transcript-level quantification and fusion detection can also be compared on our dataset. Also, we did not test the performance of tools at the pipeline-level, where different transcript count matrices based on detected isoforms could be used as input for DTE and DTU analysis. Furthermore, we did not include long-read data generated by different sequencing protocols and technology platforms. Currently, getting ∼40 million reads per sample is still prohibitively expensive for PacBio, ruling out a fair comparison between long-read sequencing platforms and the DTE/DTU analyses. The use of established cell lines and spike-ins makes it relatively easy to extend this study to include from new technologies in the future. We hope that our data and analysis can guide researchers towards better analysis of long-read RNA-seq data.

## Methods

### Cell culture, mRNA extraction and study design

Two human lung adenocarcinoma cell lines (NCI-H1975 and HCC827) from 2-4 of passages were retrieved from the American Type Culture Collection (ATCC) (https://www.atcc.org/) and cultured on 3 separate occasions in Roswell Park Memorial Institute (RPMI) 1640 medium with 10% fetal calf serum and 1% penicillin-streptomycin. The cells were grown independently at 37°*C* with 5% carbon dioxide until near 100% confluency. Cells were dissociated into single cell suspensions in FACS buffer, centrifuged and frozen at -80°*C*. The mRNA was extracted using a Qiagen RNA miniprep kit and purified using the NEBNext^®^ Poly(A) mRNA Magnetic Isolation Module (E7490). Synthetic ‘sequin’ RNA standards were then added to the cell lines – NCI-H1975 with sequins mix A, and HCC827 with sequins mix B (Figure 1a). These samples included 76 synthetic genes, each with 1-4 transcripts consisting of 1-36 exons. Libraries were then constructed for processing on 2 different platforms – ONT PCR cDNA sequencing on PromethION platform for long reads, and Illumina RNA-seq on NextSeq 500 platform for short reads.

### RNA sequencing and data preprocessing

#### Long reads

ONT cDNA libraries were constructed with SQK-PCS109 cDNA-PCR sequencing and SQK-PBK004 PCR Barcoding kits using the supplied protocol. Triplicate libraries of each mix were constructed using 1.5ng as input for cDNA synthesis. Samples were barcoded 1 to 6 using supplied PCR barcodes. Transcripts were amplified by 14 cycles of PCR with a 2-minute-30-second extension time. Pooled barcoded libraries were sequenced on 5 PromethION R9 flow cells and sequenced using the ONT PromethION platform. The fast5 files were base-called by *Guppy* version 3.2.8, available to ONT customers via the community site (https://community.nanoporetech.com/).

#### Short reads

Illumina mRNA libraries were constructed with the NEBNext^®^ Ultra II Directional RNA Library Prep Kit. Fragmentation was carried out for 15 minutes at 94°*C*, and final amplification with 11 cycles of PCR and libraries were sequenced in 75PE on NextSeq 500. Base calling and quality scoring were conducted using *Real-Time Analysis* onboard software version 2.4.6 (Illumina). The fastq files were generated and demultiplexed using *bcl2fastq* conversion software version 2.15.0.4 (Illumina).

### *In silico* mixture

Long and short reads from NCI-H1975 (000) and HCC827 (100) samples were randomly subsampled and mixed separately in 3 different ratios: 75:25, 50:50 and 25:75. To precisely control the mixing ratios and obtain equal library sizes for each sample, 7.5 million (25%), 15 million (50%) and 22.5 million (50%) reads were extracted randomly from each sample using *seqtk* (https://github.com/lh3/seqtk) using a fixed random seed (100). Each NCI-H1975 sample was paired with an HCC827 sample to create *in-silico* mixture samples from their subsampled reads – 30 million reads in total for each *in-silico* mixture sample, with 3 replicates in each mixture group (Figure 1a).

### *In vitro* mixture

To validate the in silico mixture for long-read PCR-cDNA sequencing, we mixed the NCI-H1975+sequins mixA replicate 1 RNA with the HCC827+sequins mixB replicate 1 RNA from above, in the 5 following mass ratios: 100:0, 75:25, 50:50, 25:75 and 0:100. 1 ng of each mix was then used for PCR-cDNA library preparation as before, with these two differences: 16 cycles of PCR (rather than 14), and a 0.8X final bead clean up (rather than 0.6X). 65 ng of the final library were loaded onto 1 FLO-PRO002 PromethION flow cell (R9.4.1). Reads were basecalled with Guppy 6.2.1 in ‘sup’ accuracy (`dna_r9.4.1_450bps_sup_prom.cfg` odel), disabling trimming and enabling read splitting.

### Downsampling analysis

To produce data with different library sizes, fixed numbers (1 million, 3 million, 5 million, 10 million, 15 million and 20 million) of long and short reads were extracted randomly from each sample using *seqtk* using a fixed random seed (100).

### Long-read isoform detection

We used six different tools to perform isoform detection and quantification of the long-read data: *bambu, Cupcake* (https://github.com/Magdoll/cDNA_Cupcake), *FLAIR*[21], *FLAMES, StringTie2*[23], and *TALON*[24]. Default parameters were used to run *bambu* (version 1.0.2 https://github.com/GoekeLab/bambu), *Cupcake* (version 25.2.0), *FLAMES* (version 0.1.0 (https://github.com/LuyiTian/FLAMES) and *TALON* (version 5.0.0). *Cupcake*’s script to collapse redundant isoforms using a reference genome was run with default settings on a sorted BAM file, to obtain a collapsed GFF file containing the unique transcripts in our ONT sequences. Prior to running *TALON*, we ran *TranscriptClean*[36] (version 2.0.2) using default parameters to perform reference-based error correction.

*FLAIR* (version 1.5.0) was run with optional arguments: *-v1*.*3* to specify *samtools* version 1.3+ in the *align* step; *–nvrna* in the *correct* step to keep read strands consistent after correction; and *–trust ends* to indicate long reads with minimal fragmentation during the *collapse* and *quantify* steps. *StringTie2* (version 2.1.5) was run using sorted BAM files for each sample, with the arguments *-L* to indicate long-read settings and *-G* to specify a reference annotation. BAM files were sorted using *samtools*[37, 38] (version 1.7). The GTF files from each *StringTie2* run were combined to generate a non-redundant set of isoforms using the transcript merge usage mode.

To compare the isoforms detected by all six tools, R/Bioconductor package *Repitools*[39–41] (version 1.40.0) was used to first create two-kilobase bins along the genome, and *IRanges*[42] (version 2.28.0) was then used to find overlaps between isoform coordinates and each bin. To calculate short-read splicing junction coverage and classify transcripts found by each tool into structural categories, the *SQANTI3* [43] (version 5.0) quality control protocol was additionally run with parameter *–short reads* specifying Illumina FASTQ files.

A downsampled dataset containing 10% of reads was generated with *seqtk*. The isoform detection tools were first run on this dataset with the same parameters and annotation GTF. Isoforms identified in the full and in the downsampled datasets were compared using *GffCompare*[44] (version 0.12.6) against the reference annotation file provided with the *-r* argument. An isoform detected in both datasets and in the reference annotation was a complete exact match if given the transcript class code ’= ’ in the output tracking file. If an isoform was only detected in the first or second query GTF, it was considered to be unique to the full or downsampled dataset, respectively.

The isoform detection tools were additionally run on the downsampled dataset using a modified annotation GTF for the sequins recovery analysis. To create the modified GTF, a random sample of 40 isoforms deriving from different transcripts was generated from the 120 sequin isoforms in the reference annotation. The distributions of isoform lengths and abundance levels for isoforms that were retained versus those that were removed showed no systematic differences (Supplementary Figure S16). For *Cupcake*, the *SQANTI3* quality control protocol was further required in order to annotate transcripts and identify sequin-related annotations in this analysis.

### Transcript-level quantification of detected isoforms

#### Long reads

Transcript-level counts generated by *bambu, FLAIR, FLAMES* and *TALON* were used directly in downstream analysis. *Cupcake* and *StringTie2* required some extra steps: transcript assemblies were created using annotation GTF files by *Cufflinks*[45] (version 2.1.1); and *Salmon*[46] (version 1.5.2) was used to assign long reads to individual isoforms using its ‘mapping-based mode’.

#### Short reads

Transcript assemblies generated by *FLAIR* and *FLAMES* were used directly in downstream analysis. For *bambu, Cupcake, StringTie2*, and *TALON*, transcript assemblies were created from their GTF files. *Salmon*[46] (version 1.5.2) was used to assign short reads to individual isoforms using its ‘mapping-based mode’.

### Read count processing

Transcripts were removed from both long- and short-read datasets if they had fewer than 10 counts in the long-read data.

#### Long reads

Counts were transformed into log_2_CPM values using the *cpm* function from R/Bioconductor package *edgeR*[19] (version 3.34.0) with parameter *log=TRUE*.

#### Short reads

Counts were transformed into log_2_TPM values (transcripts per million) using a prior count of 0.5 and transcript lengths generated by *Salmon* from the counting process.

### Transcriptomic alignment and transcript-level quantification against reference annotation

#### Long reads

Reads were mapped to the GENCODE Human Release 33 transcript sequences and RNA sequin transcript sequences using *minimap2* [47] (version 2.17), with arguments *-ax map-ont –sam-hit-only*. Secondary alignments were removed from BAM files using *Samtools*[37] (version 1.14). Then, we ran the ‘alignment-based mode’ in *Salmon*[46] (version 1.4.0) to obtain transcript-level quantification, where 100 bootstrap samples were computed to estimate technical variance.

#### Short reads

Transcript-level quantification was carried out by *Salmon* (version 1.4.0) on its ‘mapping-based mode’; flag *–validateMappings* was used for a selective alignment strategy, and 100 bootstrap samples were computed to estimate technical variance. For quality control purposes, reads were also mapped to the the GENCODE Human Release 33 genome and RNA sequin decoy chromosome using *STAR*[48] (version 2.7.9a), with the argument *–quantMode TranscriptomeSAM* to produce alignments translated to transcript coordinates.

Coverage fractions of transcripts by individual reads were calculated using R/Bioconductor package *GenomicAlignments*[42] (version 1.28.0) as described by Soneson *et al*.[9]. Gene body read coverage was calculated using the python package *RSeQC* [49] version 5.0.1

We annotated transcripts and gene biotypes using R/Bioconductor package *AnnotationHub* (https://bioconductor.org/packages/release/bioc/html/AnnotationHub.html version 3.2.2) to retrieve Ensembl based human annotation version 104. The function *catchSalmon* from R/Bioconductor package *edgeR*[19] (version 3.34.0) was used to read transcript-level counts and estimate transcriptome mapping ambiguity (over-dispersion coefficient) for each transcript using the bootstrap samples generated by *Salmon*. Specifically, *catchSalmon* estimates the extra variation that is introduced during read-transcript assignment over that of the baseline Poisson variation that occurs at the cDNA sampling step. The Spearman correlation between long-read CPM and short-read TPM were calculated using the *R* function *cor* with argument *method=“spearman”*. The linear relationship of sequin transcript quantification with annotated abundance was calculated by fitting a linear model using the *R* function *lm*, and the 95% confidence intervals of R-squared were calculated using the function *ci rsquared* from the *R* package *confintr*. The estimated number of transcript counts were loaded into R using R package *tximport* [50] (version 1.20.0) for differential transcript expression and differential transcript usage analysis. For each two-group comparison, we subset the data for the two groups of interest.

### Differential transcript expression analysis

We used five different tools to perform differential transcript expression analysis: *limma*[30] version 3.48.1, *edgeR*[19] version 3.34.0, *DESeq2* [28] version 1.32.0, *EBSeq* [29] version 1.32.0 and *NOISeq* [31] version 2.36.0. We used the methods recommended by each tool to perform low-count gene filtering and data normalization, and chose a representative pipeline from each tool for the DGE analysis as described in their vignettes.

When using *limma* and *edgeR*, lowly expressed genes were removed using the *filterByExpr* function from *edgeR* using default arguments. The read counts were normalized using the trimmed mean of M-values (TMM) method[51]. The *limma-voom* pipeline[52, 53] and *edgeR* quasi-likelihood pipeline[54] were used to perform differential transcript expression analysis.

When using *DESeq2*, transcripts expressed with a count of 10 or higher in at least 3 samples were kept for downstream analysis. The *DESeq* function was used to estimate size factors using the median-by-ratio method[55], estimate dispersion, fit the Negative Binomial generalized linear model and perform a Wald test to calculate *p*-values for DE.

The median-by-ratio normalization method was also used in the *EBSeq* pipeline, followed by the function *EBTest* to use the empirical Bayes hierarchical model to detect differentially expressed genes. The arguments *maxround=5* was used to run 5 iterations. We extracted the posterior probabilities of being equally expressed (PPEE) for each gene and treated genes with PPEE less than 0.05 as differentially expressed genes with false discovery rate controlled at 5%.

Low count genes were filtered out using the *filtered*.*data* function in the package *NOISeq* based on CPM values using default threshold. The *noiseqbio* function was used to compute differential gene expression between conditions, where the TMM method was used to normalize the read counts.

### Differential transcript usage analysis

We used five different methods to perform differential transcript usage analysis: *DRIMSeq* version 1.20.0[33], *DEXSeq* version 1.38.0[35], the *diffSplice* function in *limma* version 3.48.1[30], the *diffSpliceDGE* function in *edgeR* version 3.34.0[19], and *satuRn* version 1.1.1[34]. *DRIMSeq* was designed for DTU analysis with transcript-level counts and gives adjusted *p*-values at both the gene- and transcript-level. *DEXSeq, limma-diffSplice* and *edgeR-diffSplice* were designed to take exon-level short-read counts to analyze differential exon usage. We instead used transcript-level counts as input for these methods to allow DTU analysis as carried recommended by Love *et al*.[32].

Lowly expressed transcripts were filtered out using the *dmFilter* function in the *DRIMSeq* [33] package. Transcripts with 10 or more counts in at least 3 samples were kept. Genes in every sample were required to have an associated gene count (obtained by summing counts across all transcripts for a given gene) of 10 or more.

*DEXSeq* returns raw and adjusted *p*-value at the “exon”-level (which should be read as transcript-level for our analysis), and the function *perGeneQValue* computes gene-level adjusted *p*-value by summarizing per-”exon” *p*-value. For *limma-diffSplice* and *edgeR-diffSplice*, we used *t* -tests to calculate “exon”-level (which should be read as transcript-level for our analysis) adjusted *p*-values, and Simes adjustment for gene-level adjusted *p*-values. *satuRn* was designed for DTU analysis and calculates raw and adjusted *p*-values for transcripts. *satuRn* also uses empirical Bayes methods to control the false positive rate, but we still used regular FDR values due to power issues. The *perGeneQValueExact* method from *DEXSeq* was performed to calculate the per-gene adjusted *p*-value as carried out in the *satuRn* package vignette.

## Data Availability

RNA-seq data are available from Gene Expression Omnibus (GEO) under accession numbers GSE172421 (main benchmarking dataset)and GSE227000 (lab-based mixture of replicate 1).

## Software for Bechmarking Analysis

All analyses for DTE, DTU and comparisons of methods performance were run in *R* version 4.1.0[56]. Results were visualized using *ggplot2* version 3.3.5[57] and *UpSetR* version 1.4.0[58].

## Code Availability

Code used to perform these analyses and generate the figures are available from https://github.com/XueyiDong/LongReadBenchmark.

## Acknowledgements

We thank Clare Weeden and Marie-Liesse Asselin-Labat for providing the cell lines used in this study.

## Author Contributions

X.D. designed the study, conducted data analysis, generated the figures and wrote the manuscript with input from all authors. M.R.M.D conducted data analysis, generated figures and wrote the manuscript. Q.G. and M.E.R. designed the study. Q.G., L.T., J.S.J and R.B. generated benchmarking data. P.L.B devised analysis methods. Y.C., G.K.S, S.L.A., C.W.L and M.E.R. supervised the research. All authors read and approved the final manuscript.

