## Supplementary Materials for "Benchmarking long-read RNA-sequencing analysis tools using *in silico* mixtures"


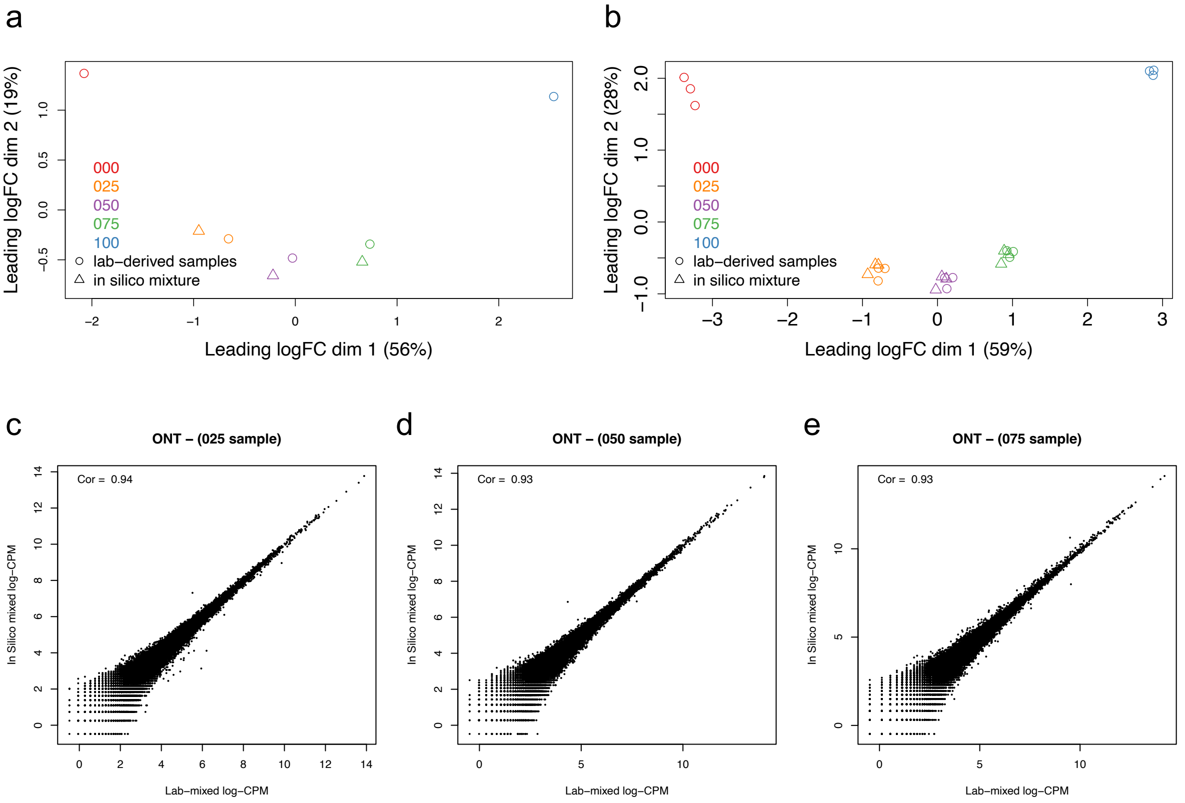


**Figure S1. Pilot study comparing *in silico* mixtures to lab-derived mixtures.** (**a-b**) MDS plots showing (**a**)ONT long-read and (**b**) Illumina short-read RNA samples mixed in the lab together with the *in-silico* mixture samples. (**c-e**) Scatter plots of the log-CPM values of transcripts from lab-derived mixture samples versus *in silico* mixture samples for mix 025 (**c**), 050 (**d**) and 075 (**e**) from the ONT dataset, along with the Spearman correlation (top-left of each plot). Data for the Illumina analysis is from the RNA-seq mixture experiment of Holik *et al.* (2017) (GSE64098).

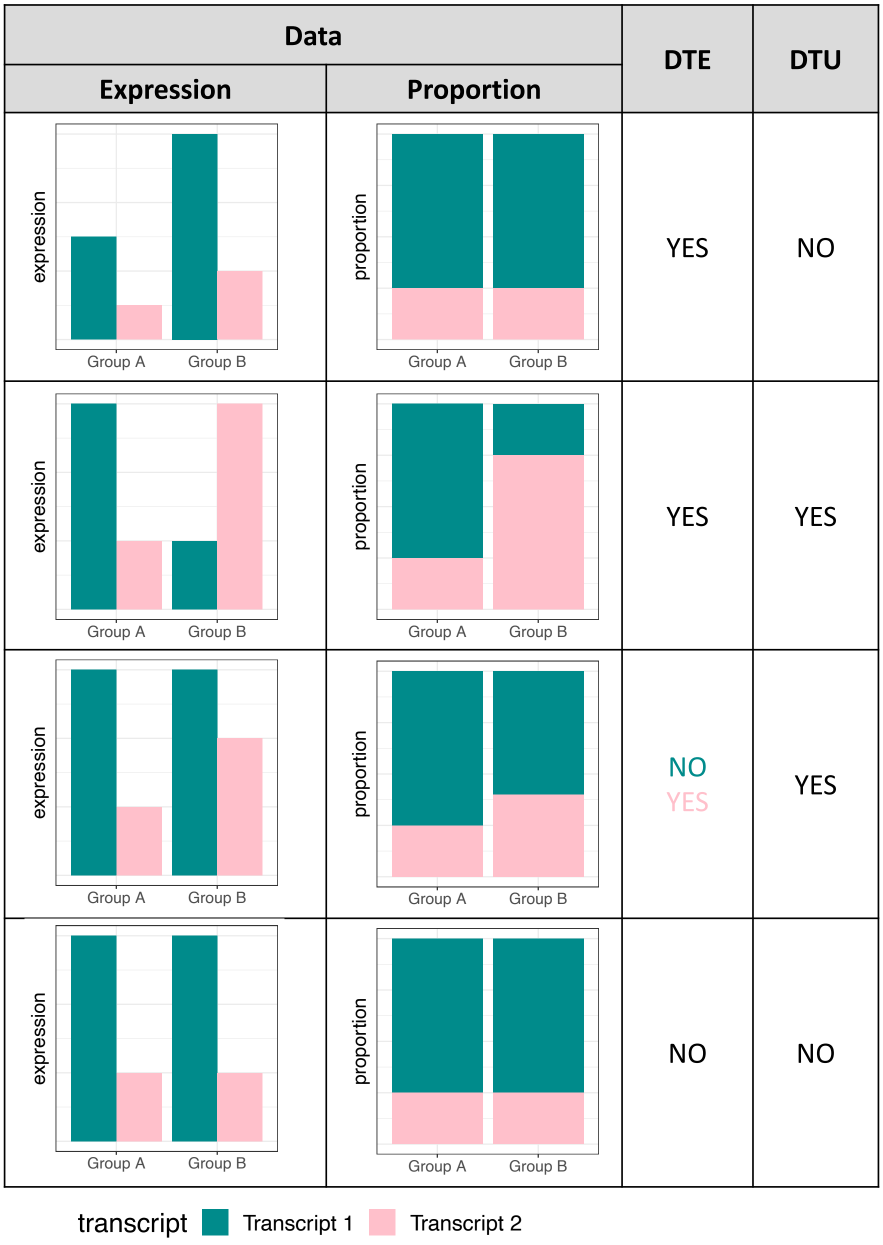


**Figure S2.** Schematic of differential transcript expression (DTE) and differential transcript usage (DTU) states for 4 genes, each with 2 transcripts. DTE analysis assesses the relative difference in expression levels between experimental groups per transcript, whereas DTU analysis assesses relative changes in proportion of expressed transcripts from a given gene between groups. Given the different hypotheses being tested in a DTE versus a DTU analysis, distinct statistical models are used in each type of analysis.

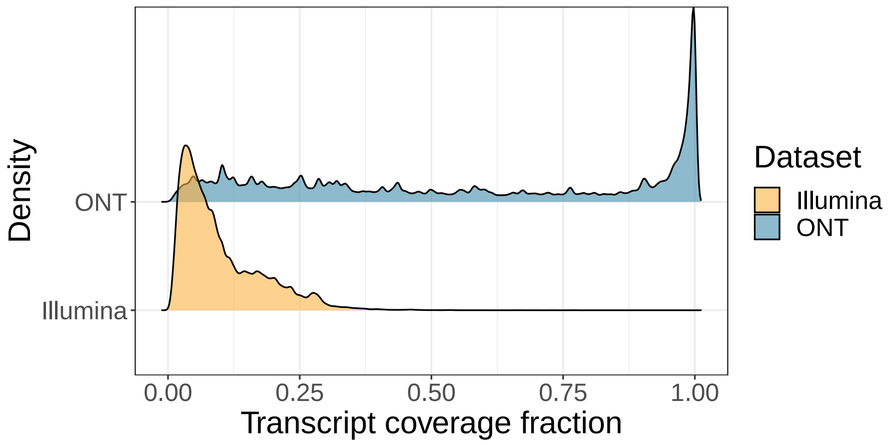


**Figure S3.** Distribution of the transcript base coverage fraction by individual ONT reads and Illumina read-pairs.

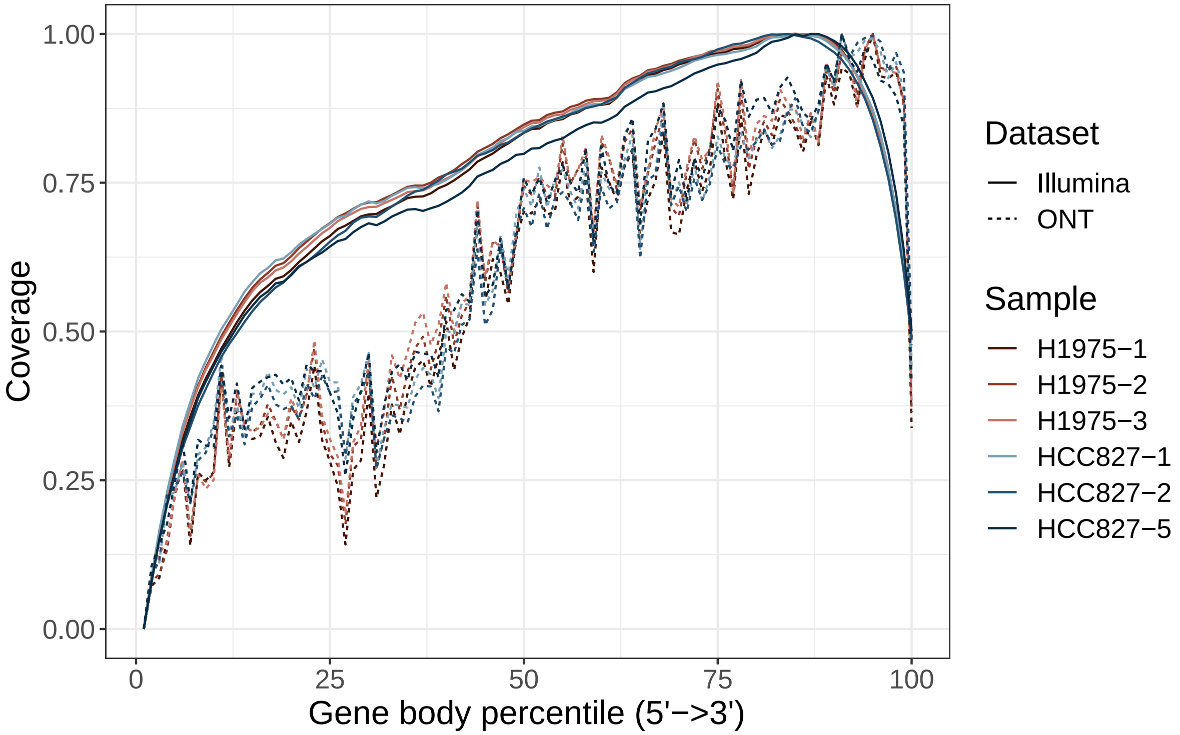


**Figure S4.** ONT (dashed lines) and Illumina (solid lines) read coverage over gene bodies (5' to 3') for each sample.

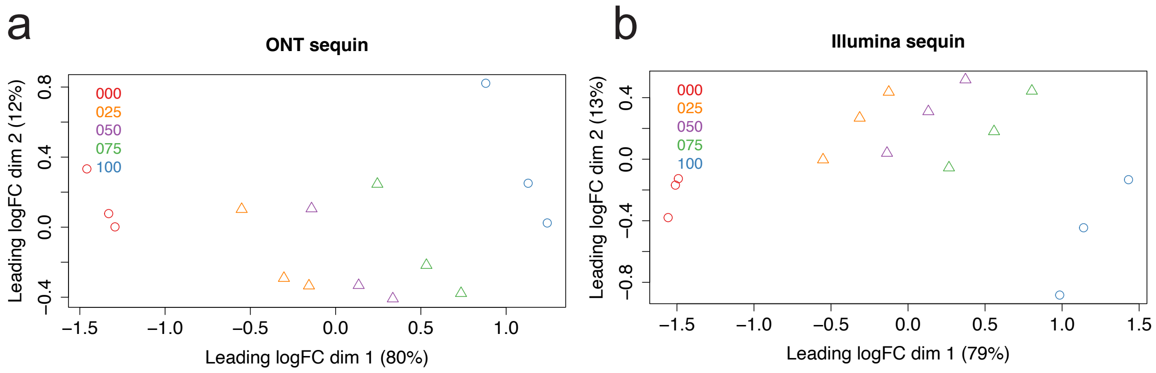


**Figure S5.** MDS plots showing the pure RNA samples and *in-silico* mixture samples based on sequins transcript-level logCPM for the ONT (**a**) and Illumina (**b**) datasets.

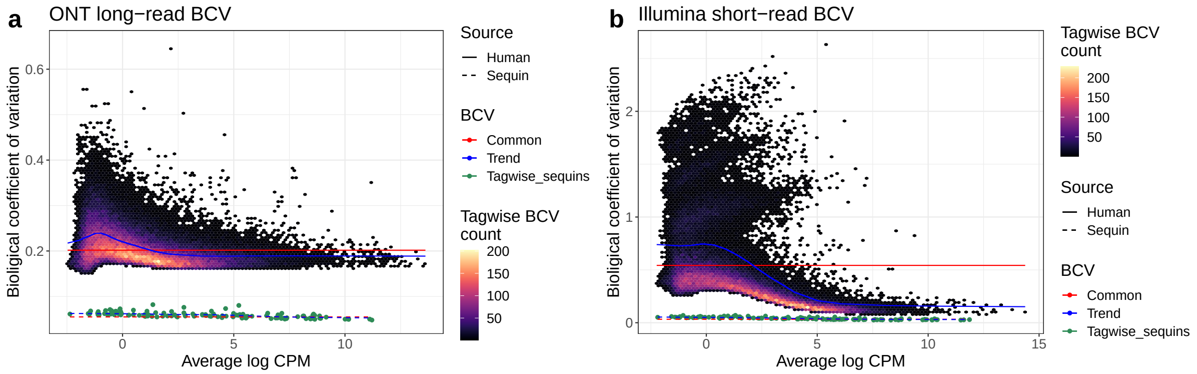


**Figure S6.** The biological coefficient of variation (BCV) against the average logCPM for each human (hexagonal 2D density and solid lines) and sequin (green points and dashed lines) transcript from the ONT (**a**) and Illumina (**b**) datasets.

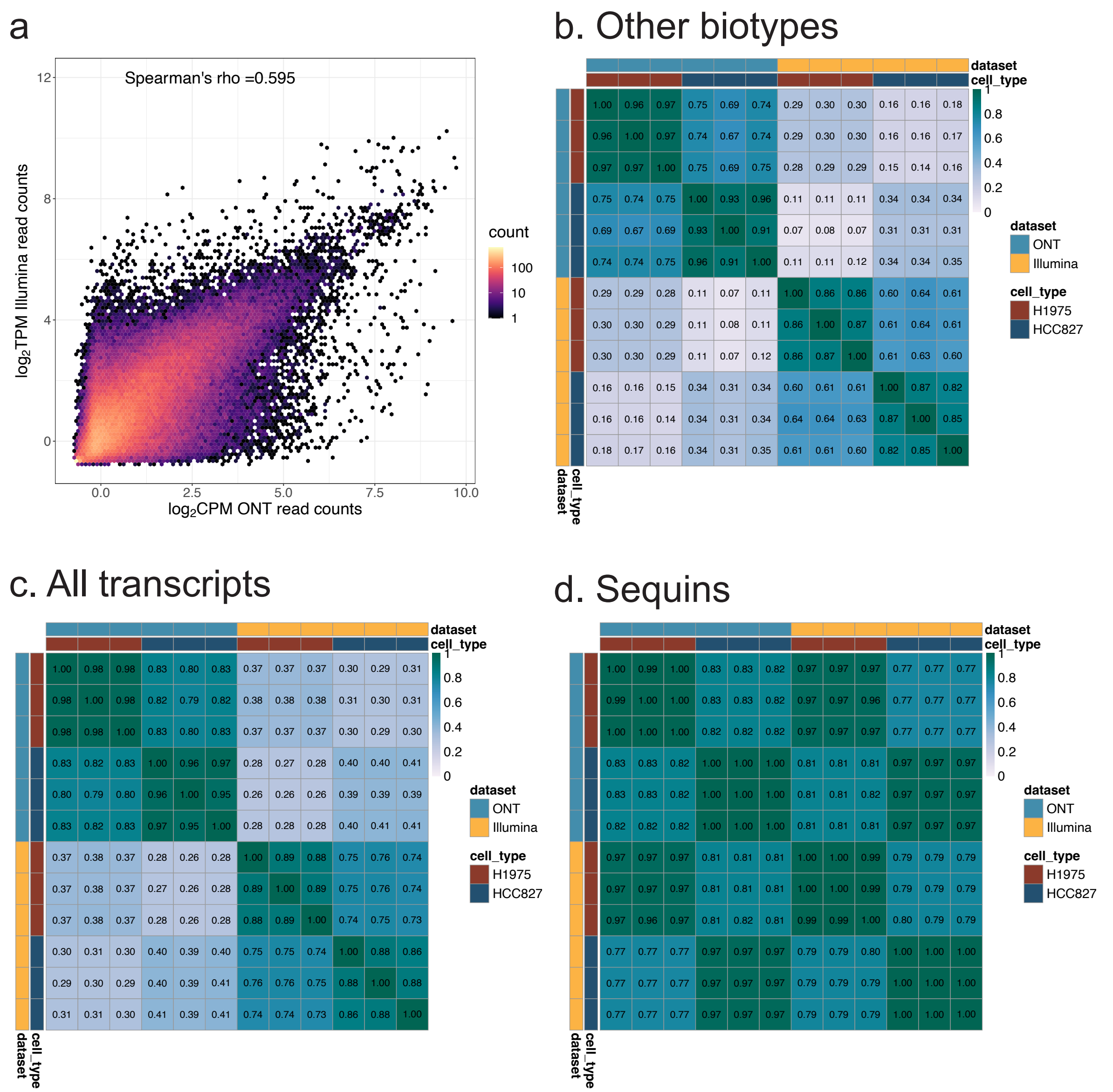


**Figure S7. Correlation of counts from different samples and/or sequencing platforms.** (**a**) A hexagonal 2D density plot showing the correlation between transcript-level quantification from the ONT ($log_2$CPM) and Illumina ($log_2$TPM) data. (**b-d**) Heatmaps showing the Spearman correlation of transcript-level CPM (ONT) and TPM (Illumina) across pure RNA samples for (**b**) human transcripts from all biotypes except protein coding and lncRNA, (**c**) all transcripts and (**d**) sequin transcripts.

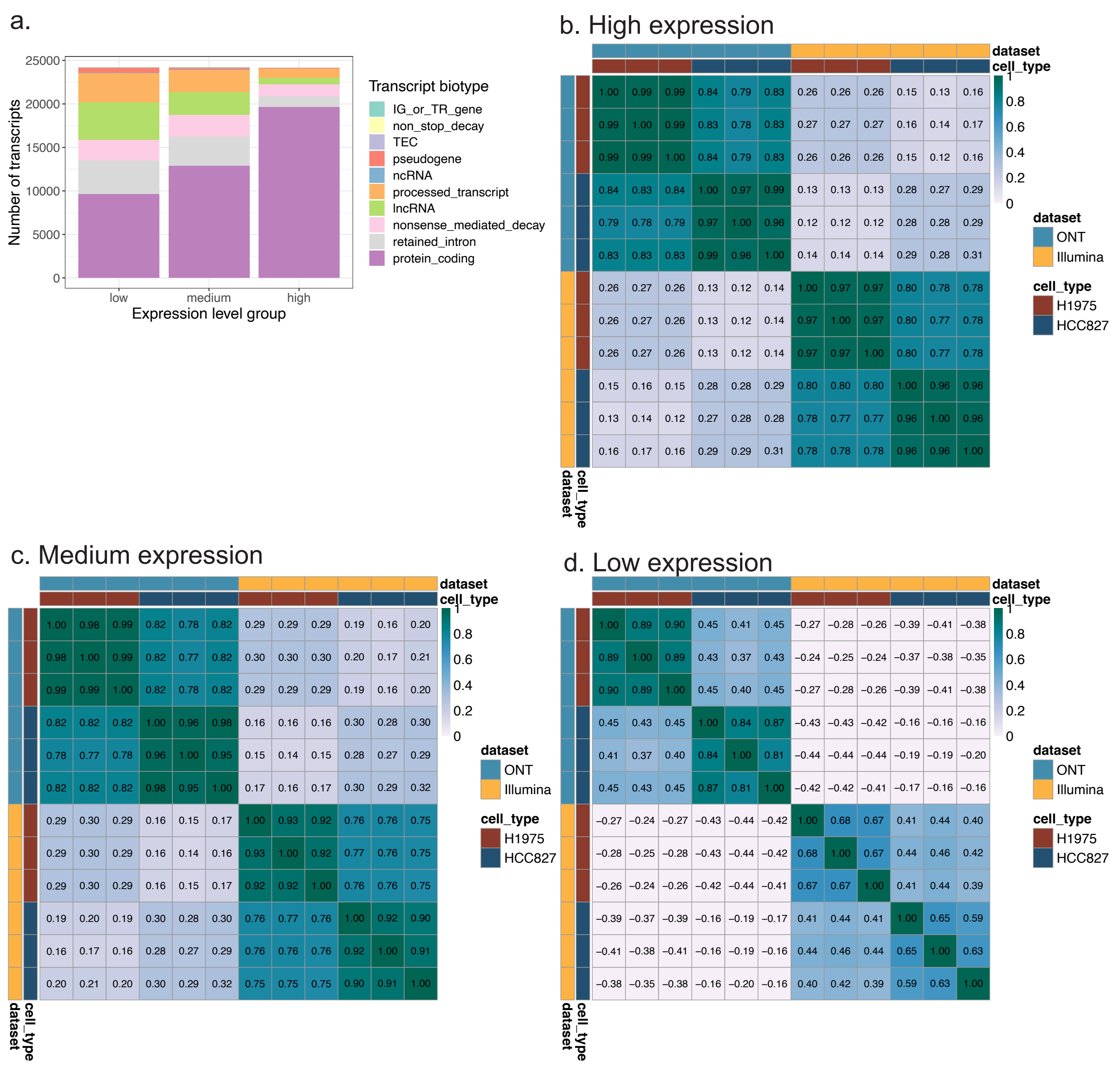


**Figure S8. Correlation of counts from different samples and/or sequencing platforms, stratified by transcript expression level.**(**a**) The number of transcripts from each biotype in each transcript expression level group, stratified by the bottom 1/3 (low) middle 1/3 (medium) and top 1/3 (high) quantiles of the total counts across all samples in both ONT and Illumina datasets. (**b-d**) Heatmaps showing the Spearman correlation of transcript-level CPM (ONT) and TPM (Illumina) across pure RNA samples for (**b**) high, (**c**) medium and (**d**) low expressed transcripts.

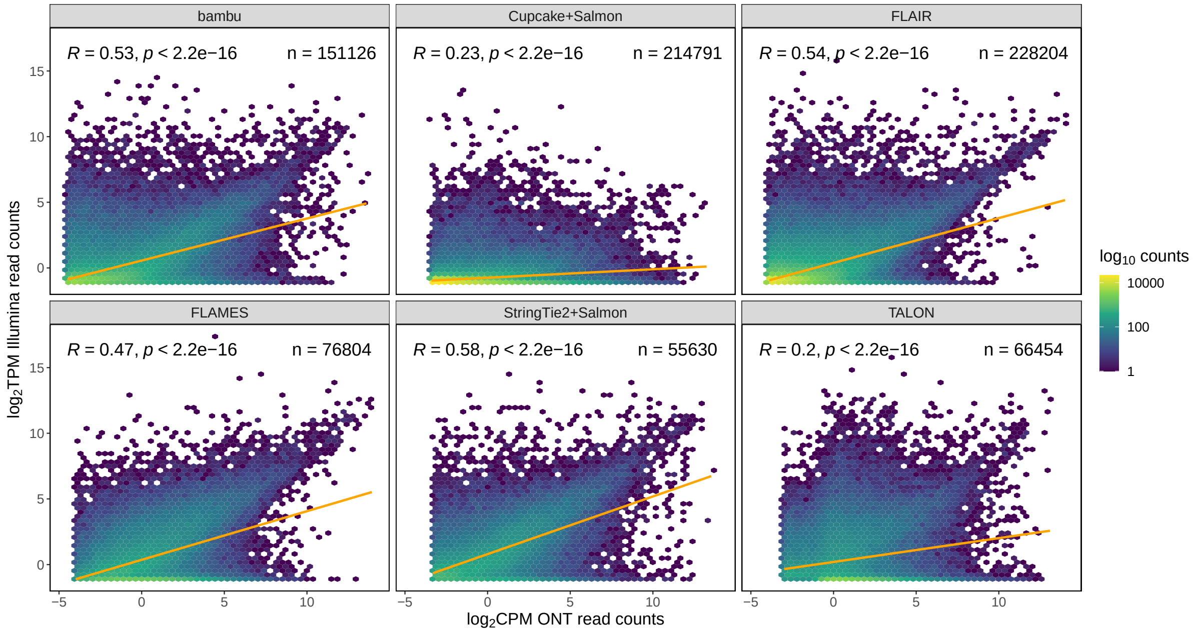


**Figure S9.** Density scatter plots showing the correlation between transcript-level ONT long-read $\log_2$CPM and Illumina short-read $\log_2$TPM (transcripts per million) quantified for each tool. A Pearson correlation coefficient and the number of features (transcripts) detected are displayed. Hexagonal bins are coloured by the read count ($\log_{10}$-scale). A regression line is highlighted in orange.

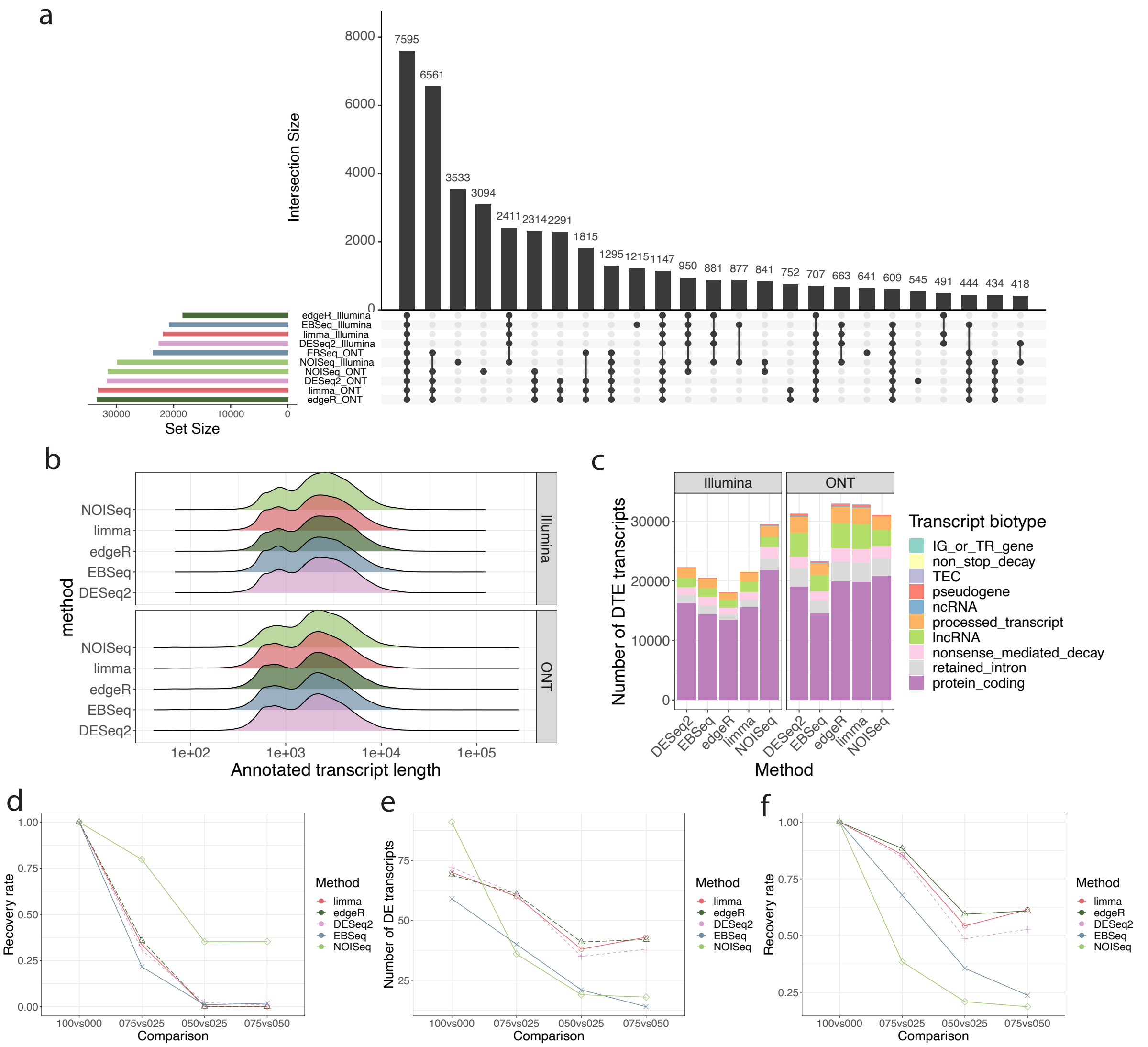


**Figure S10. Comparing differential transcript expression methods using *in silico* mixtures.** (**a**) An UpSet plot showing the 25 largest intersections of differentially expressed transcripts between the HCC827 (100) and H1975 (000) samples detected by each tool in the Illumina short-read and ONT long-read data. All transcripts are included in this analysis. (**b**) Length distribution of differentially expressed transcripts detected by each tool in the Illumina short-read (top) and ONT long-read (bottom) data. (**c**) The number of differentially expressed human transcripts stratified by biotype detected by each tool in the Illumina short-read (left) and ONT long-read (right) data. (**d**) The recovery rate of differentially expressed human transcripts detected for each method for different mixture comparisons. (**e**)The number of differentially expressed sequin transcripts detected in each comparison of the ONT data. (**f**) The recovery rate of differentially expressed sequin transcripts detected for each method for different mixture comparisons.

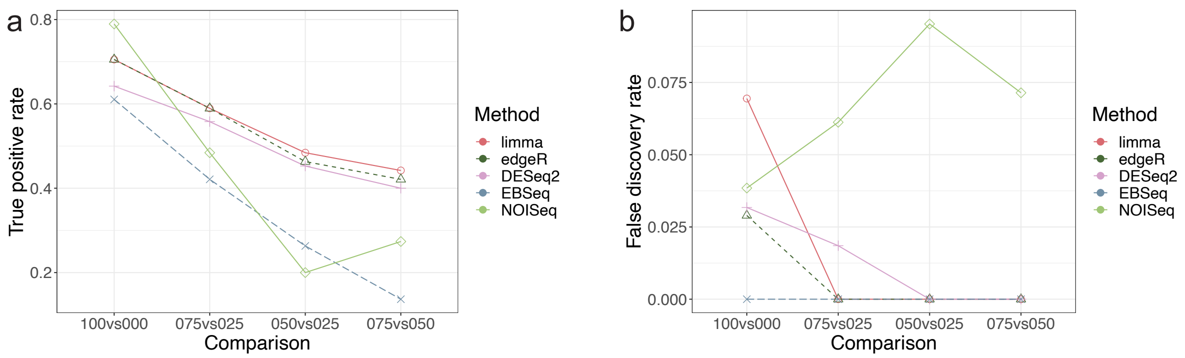


**Figure S11. Comparing differential transcript expression methods using *in silico* mixtures in the Illumina short-read data.** (**a**)The true positive rate (TPR) of different comparisons from each differential transcript expression tool in the Illumina sequins data. (**b**) The false discovery rate (FDR) of different comparisons from each differential transcript expression tool in the Illumina sequins data.

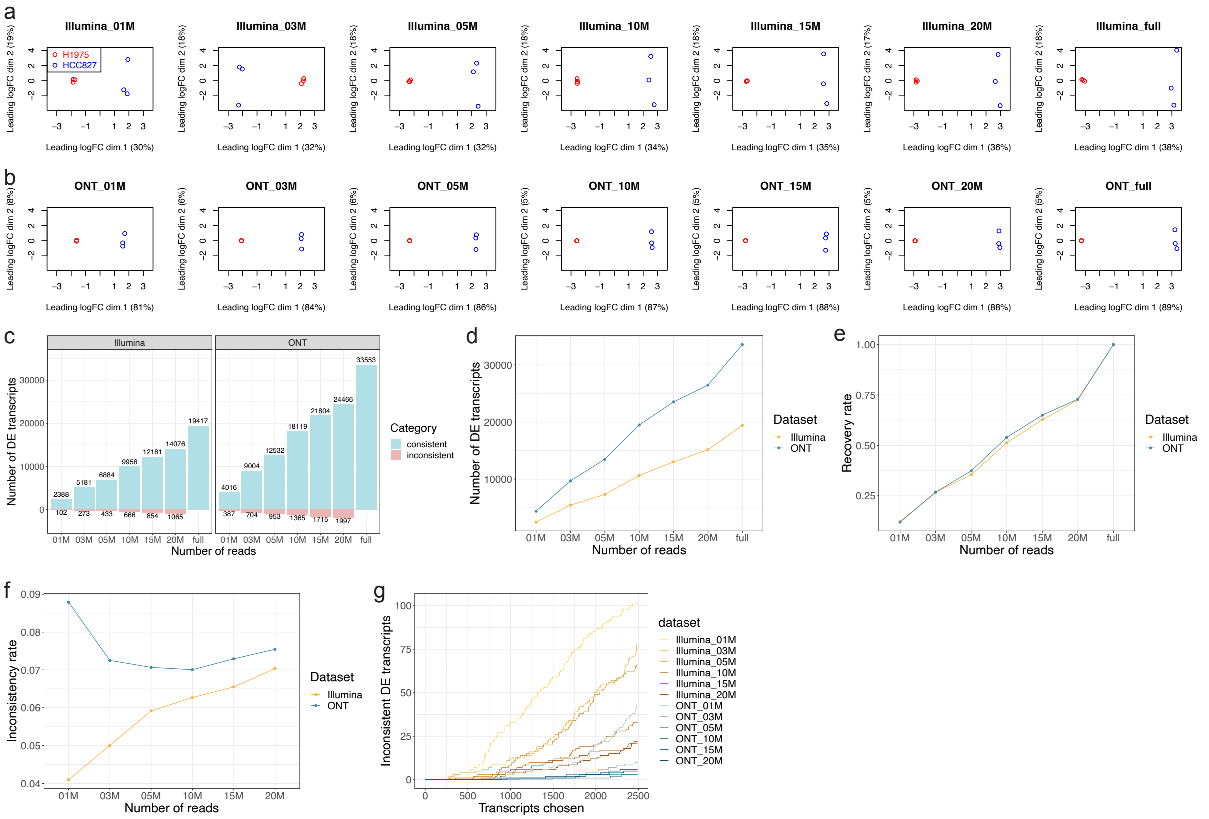


**Figure S12. Differential transcript expression analysis on dataset down-sampled to different library sizes.** (**a-b**)MDS plots showing the pure RNA samples based on human transcript-level logCPM for the down-sampled Illumina (**a**) and ONT (**b**) datasets of different library sizes. (**c**) A bar plot showing the number of consistent (blue) and inconsistent (pink) significant human DTEs detected on Illumina and ONT datasets of down-sampled to each library size. (**d**) A line plot showing the number of significant human DTEs detected on Illumina and ONT datasets downsampled to each library size. (**e**) The recovery rate of differentially expressed human transcripts detected by *edgeR* on Illumina and ONT datasets downsampled to each library size. (**f**) The Inconsistency rate of differentially expressed human transcripts detected by *edgeR* on Illumina and ONT datasets downsampled to each library size. (**g**) A line plot showing the number of inconsistent discoveries in each dataset versus the number of transcripts selected as differentially expressed.

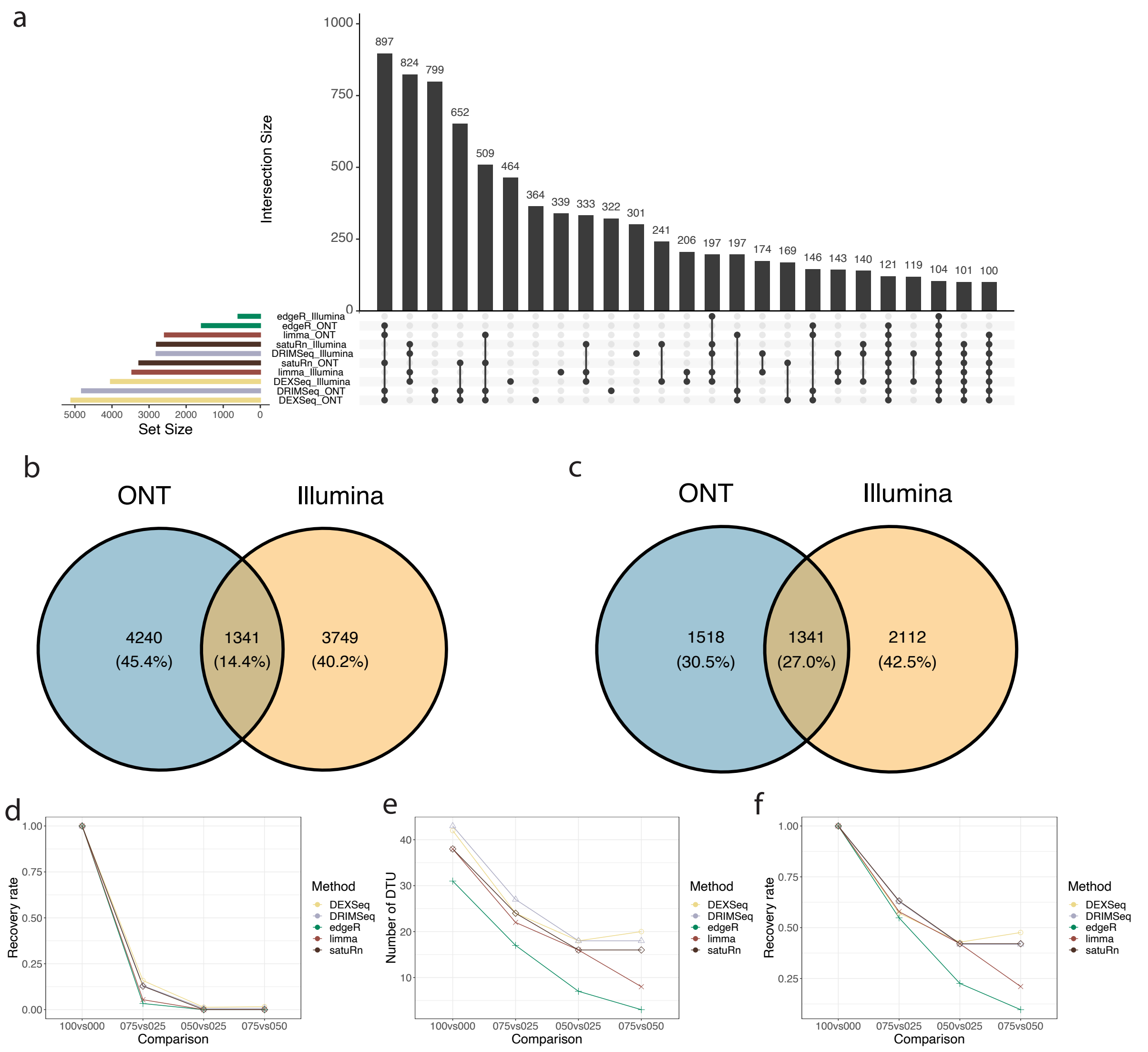


**Figure S13. Comparison of differential transcript usage (DTU) methods on *in silico* mixtures at the transcript-level.** (**a**) An UpSet plot showing the 25 largest intersections of DTU between HCC827 (100) and H1975 (000) samples detected by each tool in the Illumina short-read and ONT long-read data. All transcripts are included in this analysis. (**b**) A venn diagram showing the overlap of DTU (transcript-level analysis) detected by any method on the ONT and Illumina datasets for any transcript (i.e. those recovered in any analysis). (**c**) A venn diagram showing the overlap of DTU (transcript-level analysis) detected by any method on the ONT and Illumina datasets for the common set of transcripts tested by all methods on both the Illumina and ONT datasets. (**d**) The recovery rate of DTU for human transcripts detected by each method for different mixture comparisons. (**e**)The number of DTU sequin transcripts in each comparison of the ONT data showing the power and the ability to recover discoveries from the 100 versus 000 comparison of each method. (**f**) The recovery rate of DTU for sequin transcripts detected by each method for different mixture comparisons.

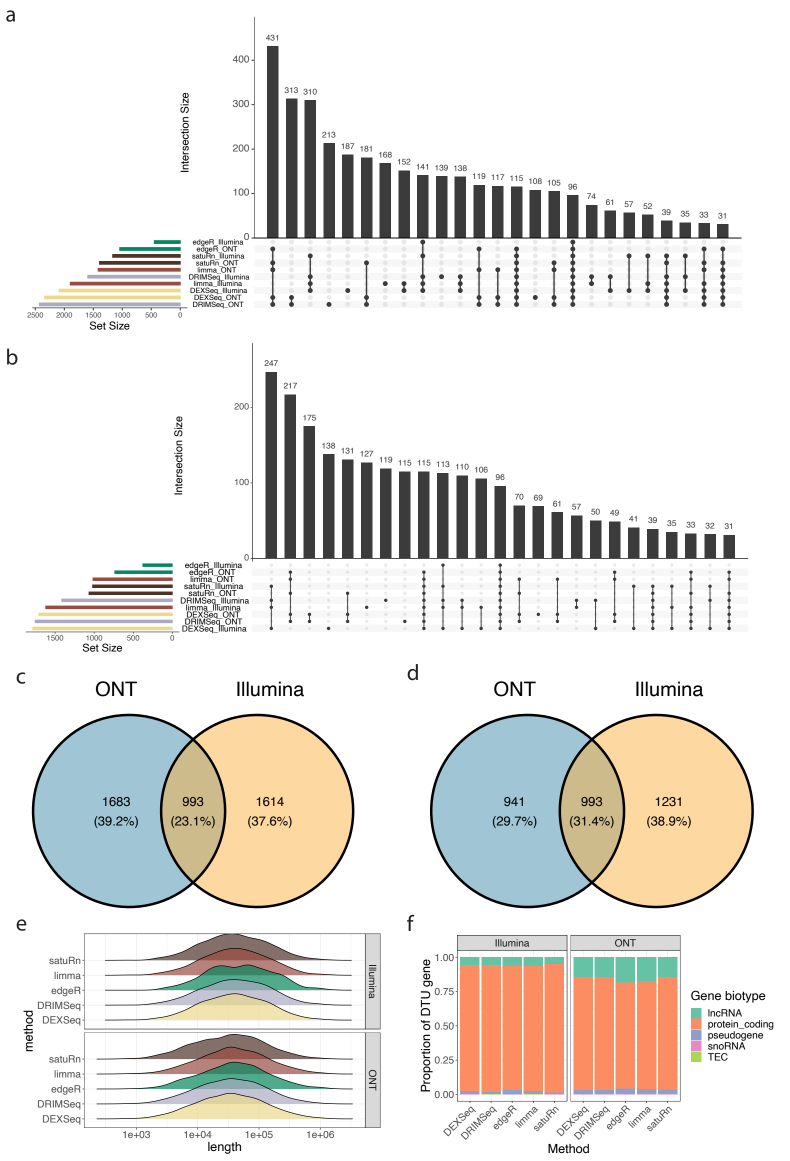


**Figure S14. Gene-level comparisons of differential transcript usage (DTU) methods on pure RNA samples.** (**a**) An UpSet plot showing the 25 largest intersections of DTU (gene-level) that contain DTU between the HCC827 (100) and H1975 (000) samples detected by each tool in the Illumina short-read and ONT long-read data for any gene (i.e. those recovered in any analysis). (**b**) An UpSet plot showing the 25 largest intersections of DTU (gene-level) that contain DTU between the HCC827 (100) and H1975 (000) samples detected by each tool in the Illumina short-read and ONT long-read data for the common set of genes tested by all methods on both the Illumina and ONT datasets. (**c**) A venn diagram showing the overlap of genes detected to have DTU by any method on the ONT and Illumina datasets for genes detected by any method. (**d**) A venn diagram showing the overlap of genes detected to have DTU by any method on the ONT and Illumina datasets for the common set of genes tested by all methods across both sequencing platforms. (**e**) Length distribution of genes with DTU detected by each tool in the Illumina short-read (top) and ONT long-read (bottom) data. (**f**) The proportion of genes detected with DTU per biotype for each tool in the Illumina short-read (left) and ONT long-read (right) data.

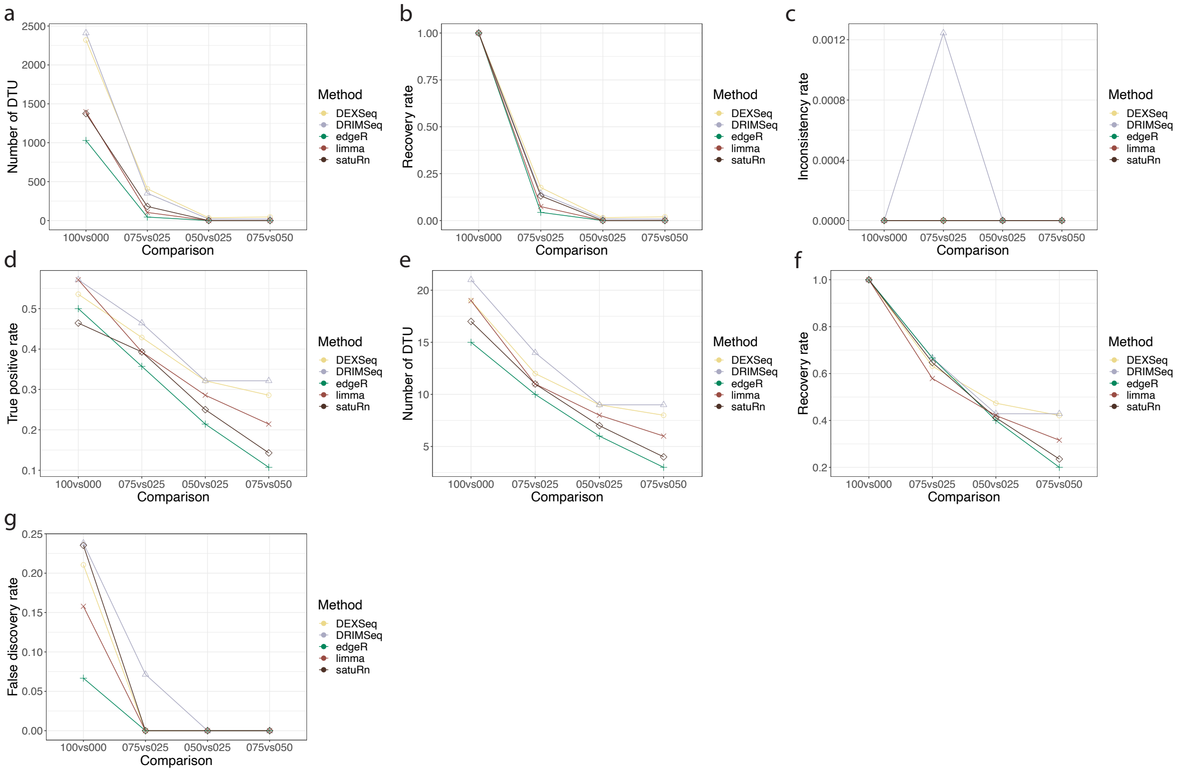


**Figure S15. Gene-level comparisons of differential transcript usage (DTU) methods using *in silico* mixtures of the ONT data.** (**a**) The number of human genes detected to have DTU in each comparison. (**b**) The recovery rate of human genes detected with DTU for each method. (**c**) The rate of human genes detected with DTU that were inconsistent with discoveries made in the 100 versus 000 comparison for each method. (**d**) The true positive rate (TPR) for different comparisons and DTU tools applied to the sequin data. (**e**)The number of sequin genes with DTU in each comparison of the ONT data showing the power and the ability to recover discoveries from the 100 versus 000 comparison of each method. (**f**) The recovery rate of sequin genes detected with DTU for different comparisons and methods. (**g**) The false discovery rate (FDR) for different comparisons and DTU tools applied to the sequin data.

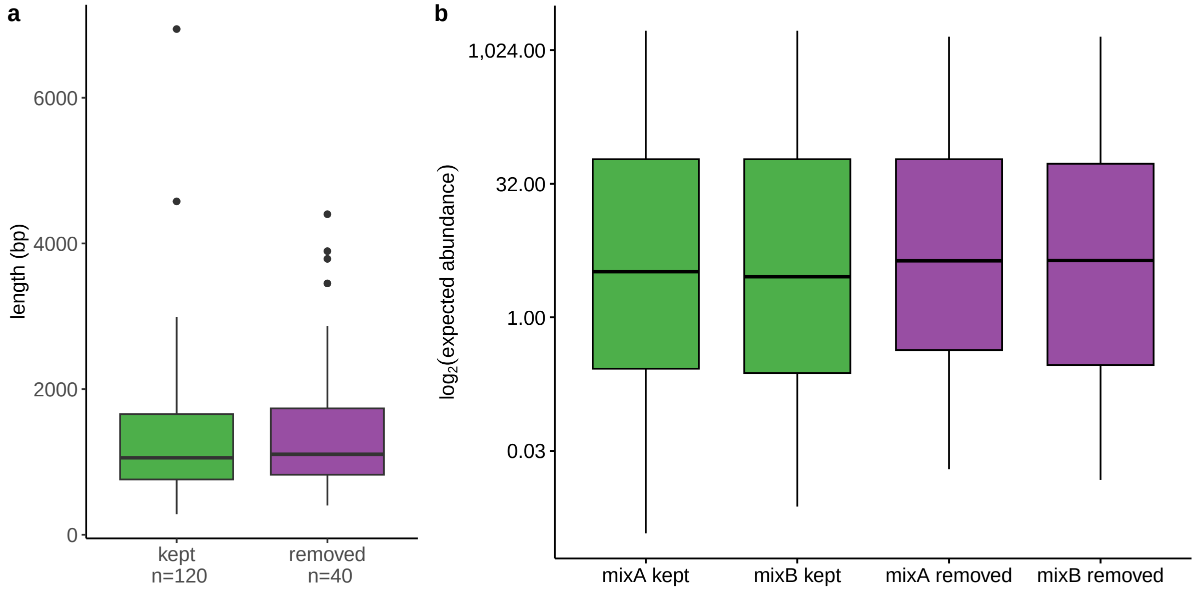


**Figure S16. Comparison of lengths and abundances for the sequin recovery analysis.** (**a**) A boxplot showing the distributions of lengths (bp) of sequin isoforms removed from or retained in the modified reference annotation. (**b**) A boxplot showing the distributions of abundance levels of the sequin isoforms on a $\log_2$-scale, coloured by removal status, and further categorised by spike-in control mix (A and B).
